# SWR1-independent association of H2A.Z to the LINC complex promotes meiotic chromosome motion

**DOI:** 10.1101/2020.06.30.181016

**Authors:** Sara González-Arranz, Jennifer M. Gardner, Zulin Yu, Neem J. Patel, Jonna Heldrich, Beatriz Santos, Jesús A. Carballo, Sue L. Jaspersen, Andreas Hochwagen, Pedro A. San-Segundo

## Abstract

The H2A.Z histone variant is deposited into chromatin by the SWR1 complex affecting multiple aspects of meiosis. Here we describe a SWR1-independent localization of H2A.Z at meiotic telomeres and the centrosome. We demonstrate that H2A.Z colocalizes and interacts with Mps3, the SUN component of the LINC complex that spans the nuclear envelope and links meiotic telomeres to the cytoskeleton promoting meiotic chromosome movement. H2A.Z also interacts with the meiosis-specific Ndj1 protein that anchors telomeres to the nuclear periphery via Mps3. Telomeric localization of H2A.Z depends on Ndj1 and the N-terminal domain of Mps3. Although telomeric attachment to the nuclear envelope is maintained in the absence of H2A.Z, the distribution of Mps3 is altered. The velocity of chromosome movement during meiotic prophase I is reduced in the *htz1Δ* mutant lacking H2A.Z, but it is unaffected in *swr1Δ* cells. We reveal that H2A.Z is an additional LINC-associated factor that contributes to promote telomere-driven chromosome motion critical for error-free gametogenesis.

## INTRODUCTION

Meiosis is a special form of cell division that lies at the heart of gametogenesis in most sexually reproducing organisms. During meiosis, a series of complex DNA and chromosome interactions culminate in the accurate segregation of a haploid complement of chromosomes to the gametes (Duro and Marston, 2015; Hunter, 2015; Keeney et al., 2014; Zickler and Kleckner, 2015). Chromatin remodeling events, including histone post-translational modifications and incorporation of histone variants, play important roles in several processes during meiotic development (Brachet et al., 2012; Crichton et al., 2014; Ontoso et al., 2014; Yamada et al., 2018a; Yamada and Ohta, 2013).

H2A.Z is a variant of the canonical H2A histone that is incorporated into chromatin by the action of the ATP-dependent SWR1 remodeling complex. SWR1 replaces an H2A-H2B dimer by H2A.Z-H2B at defined nucleosomes, preferentially in the vicinity of promoter regions (Luk et al., 2010; Raisner et al., 2005). H2A.Z participates in a number of fundamental biological processes in vegetative cells including transcription regulation, chromatin silencing, DNA damage response and chromosome segregation (Adkins et al., 2013; Billon and Cote, 2013; Morillo-Huesca et al., 2010; Weber and Henikoff, 2014). In addition, although the number of meiotic reports is scarce, the roles of H2A.Z during meiosis are also beginning to be elucidated in some model organisms. In plants and fission yeast, H2A.Z is required for initiation of meiotic recombination, although the precise event influenced by H2A.Z appears to be different in both organisms. In *Arabidopsis thaliana*, H2A.Z has been proposed to regulate meiotic double-stand break (DSB) formation and repair by its association to hotspots and by controlling the expression pattern of recombination genes, whereas in *Schizosaccharomyces pombe*, H2A.Z impacts meiotic recombination by modulating chromatin architecture and the binding of DSB-formation proteins to cohesin-rich domains (Choi et al., 2013; Qin et al., 2014; Rosa et al., 2013; Yamada et al., 2018b). H2A.Z is also required for proper meiotic development in *Saccharomyces cerevisiae*. The budding yeast *htzłΔ* mutant (lacking H2A.Z) displays slower kinetics of meiotic progression, reduced spore viability and misregulated meiotic gene expression, but meiotic recombination is not, at least drastically, affected. In addition, the meiotic checkpoint response triggered by the absence of the synaptonemal complex (SC) Zip1 protein is altered in the *htz1Δ* mutant (Gonzalez-Arranz et al., 2018). Not surprisingly, many of the meiotic functions mentioned above for H2A.Z rely on its chromatin deposition mediated by the SWR1 complex.

Curiously, the physical interaction of H2A.Z with non-chromatin components has been reported in high-throughput analyses in *S. cerevisiae* (Bommi et al., 2019; Uetz et al., 2000; Yu et al., 2008). In particular, H2A.Z interacts with the SUN domain-containing Mps3 protein (Gardner et al., 2011). SUN proteins are one of the main components of the evolutionarily conserved LINC (Linker of the Nucleoskeleton and Cytoskeleton) complex that physically connects the nuclear contents with cytoskeletal filaments. The LINC complex is composed of an ensemble of KASH-SUN proteins. The KASH proteins span the outer nuclear membrane to interact with the cytoskeleton and also interact in the perinuclear space with SUN proteins. The SUN proteins, in turn, are embedded in the inner nuclear membrane and protrude towards the nuclear inside (reviewed by (Chang et al., 2015)). LINC complexes participate in a number of cellular functions, such as nuclear positioning, centrosome dynamics and attachment to the nuclear envelope (NE), and DNA Repair (Fernandez-Alvarez et al., 2016; Friederichs et al., 2011; Lawrence et al., 2016; Lee and Burke, 2018). LINC complexes also play a fundamental role in meiotic chromosome movement in all organisms studied (reviewed by (Burke, 2018; Link and Jantsch, 2019)). In budding yeast, two KASH proteins (Mps2 and Csm4) and one SUN protein (Mps3) have been described. In mitotic cells, Mps2 localizes at the yeast centrosome-equivalent called spindle pole body (SPB) forming a non-canonical LINC complex with Mps3 (Chen et al., 2019). In contrast, the meiotically induced Csm4 protein forms LINC complexes with Mps3 along the NE. A recent report has shown that, during meiotic prophase, Mps2 also interacts with Csm4 at the NE mediating the coupling with the Myo2 microfilament motor associated to the actin cytoskeleton (Lee et al., 2020). During meiotic prophase, telomeres are anchored to the nucleoplasmic N-terminal domain of Mps3 by the mediation of the meiosis-specific Ndj1 protein. Forces generated in the cytoplasm by the actin cytoskeleton are transduced through the Myo2/Csm4-Mps3-Ndj1 axis to promote telomere-led chromosome movement (Figure 1A). Besides facilitating homolog interactions that sustain meiotic recombination and chromosome synapsis, these movements have been also proposed to be important to disengage non-homologous chromosome links (Chua and Roeder, 1997; Conrad et al., 1997; Conrad et al., 2008; Conrad et al., 2007; Kosaka et al., 2008; Koszul et al., 2008; Rao et al., 2011; Trelles-Sticken et al., 2000; Wanat et al., 2008).

**Figure 1.**
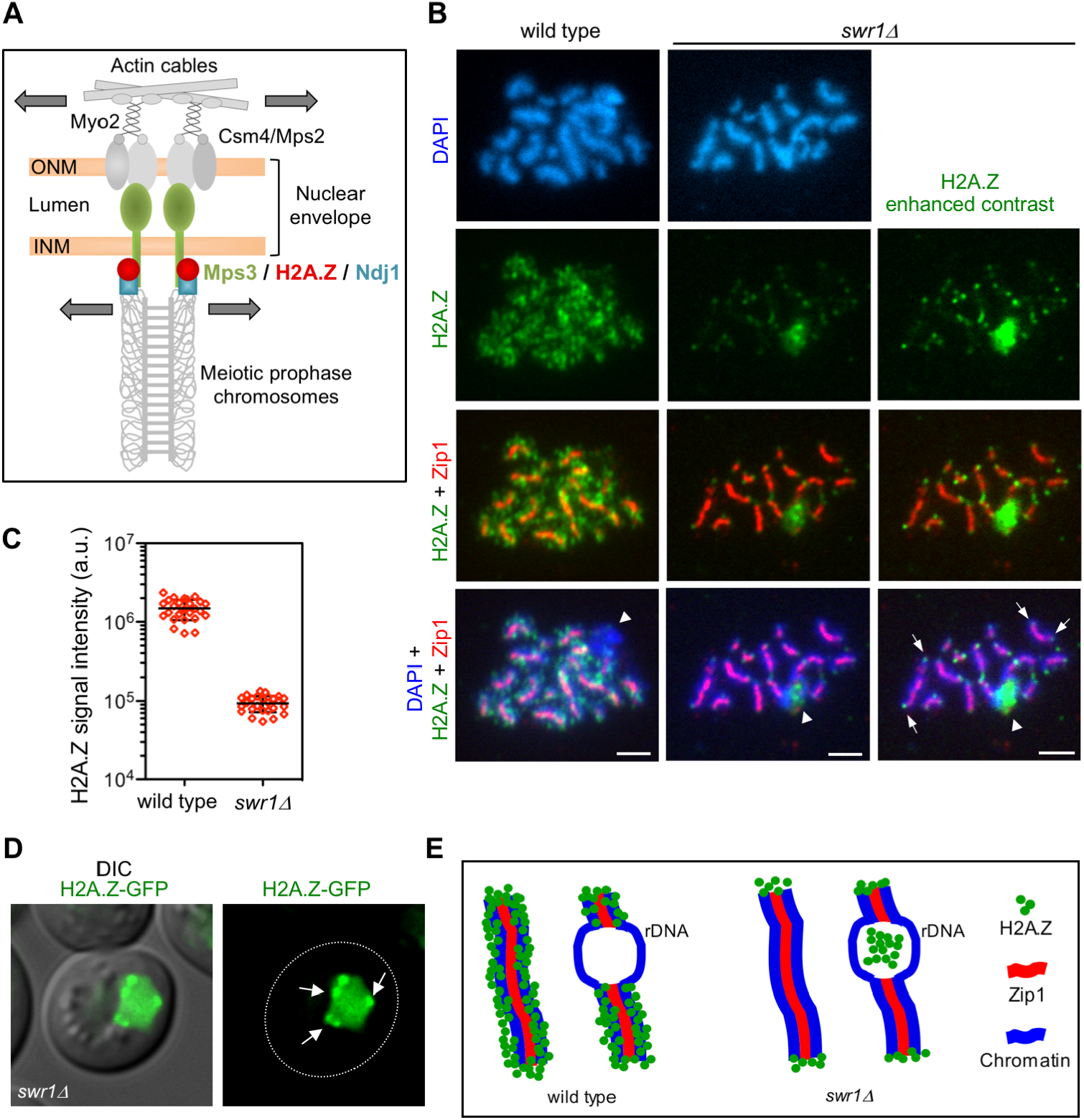
H2A.Z localizes to chromosome ends during meiotic prophase I. **(A)** Model for LINC-dependent meiotic chromosome movement. Schematic representation of the main components involved in promoting chromosome motion during meiotic prophase I in *Saccharomyces cerevisiae*, including H2A.Z as described here. Colored proteins are the focus of this work. Motion is depicted with arrows. See text for details. ONM: Outer nuclear membrane. INM: Inner nuclear membrane. **(B)** Immunofluorescence of spread pachytene chromosomes from wild type and *swr1Δ* stained with DAPI to visualize chromatin (blue), anti-GFP to detect H2A.Z (green), and anti-Zip1 to mark the SC central region (red). The H2A.Z signal in *swr1Δ* was obtained using a 4-times longer exposure time compared to the wild type. In addition, the contrast of the image shown in the rightmost column was computer-enhanced. Representative nuclei are shown. Spreads were prepared 16 h after meiotic induction. Arrows point to some telomeric H2A.Z foci. Arrowheads mark the nucleolar rDNA region devoid of Zip1. Scale bar, 2 μm. **(C)** Quantification of the total H2A.Z signal in nuclear spreads prepared as in (B). 27 and 26 nuclei from wild type and *swr1Δ*, respectively, were scored. Mean and standard deviation values are represented. **(D)** Representative images from a *swr1Δ* live cell expressing *HTZ1-GFP* 16 h after meiotic induction. Arrows point to H2A.Z foci at the nuclear periphery that stick out over the diffuse pan-nuclear H2A.Z-GFP signal. Scale bar, 2 μm. **(E)** Cartoon representing H2A.Z localization in wild-type and *swr1Δ* pachytene chromosome based on our cytological observations. Strains in (B-C) are: DP840 (*HTZ1-GFP*) and DP841 (*HTZ1-GFP swr1Δ*). Strain in (D) is DP1108 (*HTZ1-GFP swr1Δ*).

Recent studies have revealed that the telomere-associated Ndj1 protein also localizes to the SPB during meiotic prophase by interacting with Mps3. Ndj1 protein stability in combination with controlled proteolysis of Mps3 at the SPB half-bridge regulate the separation of duplicated SPBs upon meiosis I entry (Li et al., 2017; Li et al., 2015). These and other observations in different organisms support a connection between telomere and centrosome functions that coordinate NE dynamics and meiotic progression (Fernandez-Alvarez and Cooper, 2017).

Although most of the roles of the H2A.Z histone variant have been ascribed to its SWR1-dependent chromatin deposition, here we characterize in detail a SWR1-independent interaction of H2A.Z with LINC-associated components, including Mps3 and Ndj1, during meiotic prophase. We show that H2A.Z colocalizes with the SUN protein Mps3 at telomeres and demonstrate that the Mps3-H2A.Z interaction does not occur in the context of chromatin, but depends on the stable attachment of telomeres to the NE. We demonstrate that H2A.Z is an additional novel factor connected with LINC complexes during meiotic prophase that is required for proper meiotic chromosome movement (Figure 1A). We propose that at least some of the meiotic defects of the *htz1* mutant may stem from faulty processes impacted by LINC function.

## RESULTS

### H2A.Z remains at chromosome ends in the absence of SWR1

Our previous cytological studies have shown that H2A.Z extensively decorates meiotic chromatin in wild-type pachytene chromosomes, except in the rDNA region where its presence is markedly reduced (Gonzalez-Arranz et al., 2018). In the *swr1Δ* mutant, the bulk of chromatin-associated H2A.Z is lost (Gonzalez-Arranz et al., 2018), but H2A.Z foci persisted in the absence of the SWR1 complex, as seen in enhanced images (Figure 1B, 1C). Double staining with Zip1 antibodies, as a marker of synapsed chromosomes, showed that these SWR1-independent H2A.Z foci were primarily located at chromosome ends (Figure 1B); 65.4% of *swr1Δ* spread nuclei scored (n=26) displayed H2A.Z at telomeres. Consistent with this telomeric localization, *swr1Δ* live meiotic cells expressing a functional *HTZ1-GFP* fusion occasionally displayed H2A.Z spots concentrated at the nuclear periphery in addition to a diffused pan-nuclear signal (Figure 1D). Curiously, whereas H2A.Z is largely excluded from the rDNA chromatin in wild-type nuclei (88.9%; n=27 nuclei), the *swr1Δ* mutant displayed an accumulation of H2A.Z in the nucleolar area (Figure 1B, arrowhead) (50.0%; n=26 nuclei). Nevertheless, this amorphous H2A.Z mass was mainly found inside the loop defined by the unsynapsed rDNA array characteristic of pachytene chromosomes, but it did not appear to be associated with chromatin. A schematic representation of H2A.Z chromosomal localization at pachytene based on cytological observations, in both wild type and *swr1Δ*, is depicted in Figure 1E.

### Genome-wide association of H2A.Z to meiotic chromatin requires SWR1

Next, we used ChIP-seq to confirm at higher resolution the SWR1 dependency for H2A.Z binding to chromatin during meiotic prophase I. Samples from cultures of wild-type and *swr1Δ* cells expressing *HTZ1-GFP* were processed at zero and 15 hours after meiotic induction. Samples from an untagged control were also taken at the same time points. Profiles of H2A.Z distribution in all chromosomes showed no strong difference between mitotic (t=0h) and meiotic prophase cells (t=15h) and revealed that genome-wide incorporation of H2A.Z is abolished in the absence of SWR1 (Figure S1A-S1C). Consistent with previous reports in mitotic cells showing that H2A.Z is enriched at the nucleosomes flanking the transcription start site of genes (Luk et al., 2010; Raisner et al., 2005), our analysis of H2A.Z position relative to ORFs revealed that, indeed, H2A.Z was enriched at the beginning of ORFs in mitotic cells, thus validating this ChIP-seq study (Figure S1D). We also found the same situation in meiotic cells (Figure S1E). Importantly, the meta-ORF profiles of the *swr1Δ* mutant were similar to those of the untagged control implying that H2A.Z chromatin binding to gene promoters was completely abolished in the absence of SWR1 (Figure S1D, S1E). Thus, like in mitotic cells, the SWR1 complex is absolutely required for genome-wide H2A.Z chromatin deposition also during meiotic prophase I.

### H2A.Z interacts and colocalizes with Mps3 at telomeres during meiotic prophase

Previous studies in vegetative cells have described a physical interaction between H2A.Z and the Mps3 protein (Gardner et al., 2011; Morillo-Huesca et al., 2019). Moreover, a recent study also reported co-purification of H2A.Z and Mps3 from meiotic cells using mass spectrometry analysis (Bommi et al., 2019). Consistent with these observations, we found that the H2A.Z-GFP foci detected in some *swr1Δ* live meiotic cells at the nuclear periphery colocalized with Mps3-mCherry (Figure 2A, arrows). Note that there was also a peripheral zone with H2A.Z-GFP, but devoid of Mps3-mCherry (Figure 2A, arrowhead), that corresponds to the accumulation of H2A.Z observed in the vicinity of the nucleolar area in the *swr1Δ* mutant (Figure 1B, 1E; Figure S2; see also Figure 2B). We next used chromosome spreading for a more detailed analysis of H2A.Z and Mps3 colocalization. It has been shown that, despite being embedded in the inner nuclear membrane, the Mps3 protein remains associated to the telomeres in spread preparations of meiotic prophase nuclei (Conrad et al., 2007; Lee et al., 2012). We detected a significant colocalization of H2A.Z and Mps3 foci at telomeres of meiotic prophase chromosomes (Pearson’s correlation coefficient 0.741; n=27 nuclei) (Figure 2B), suggesting that H2A.Z and Mps3 also interact during meiotic prophase. To confirm the meiotic interaction between H2A.Z and Mps3 we carried out co-immunoprecipitation experiments. We found that Mps3 (tagged with 3HA) was detected in immunoprecipitates of *htz1-GFP* strains pulled down with anti-GFP antibodies (Figure 2C). Conversely, H2A.Z was present in immunoprecipitates of *MPS3-GFP* meiotic cells pulled down with anti-GFP antibodies (Figure 2D). Although the amount of H2A.Z was reduced in whole cell extracts (WCE) from the *swr1Δ* mutant, co-immunoprecipitation of H2A.Z and Mps3 occurred both in wild-type and *swr1Δ* meiotic cells (Figure 2C, 2D). Of note, immunoprecipitation of Mps3-GFP specifically brought down H2A.Z, but not the canonical histones (Figure 2D). In sum, these observations indicate that Mps3 and H2A.Z physically interact during meiotic prophase in a SWR1-independent manner. The colocalization at the end of chromosomes suggests that the Mps3-H2A.Z interaction occurs, at least, in the proximity of telomeres.

**Figure 2.**
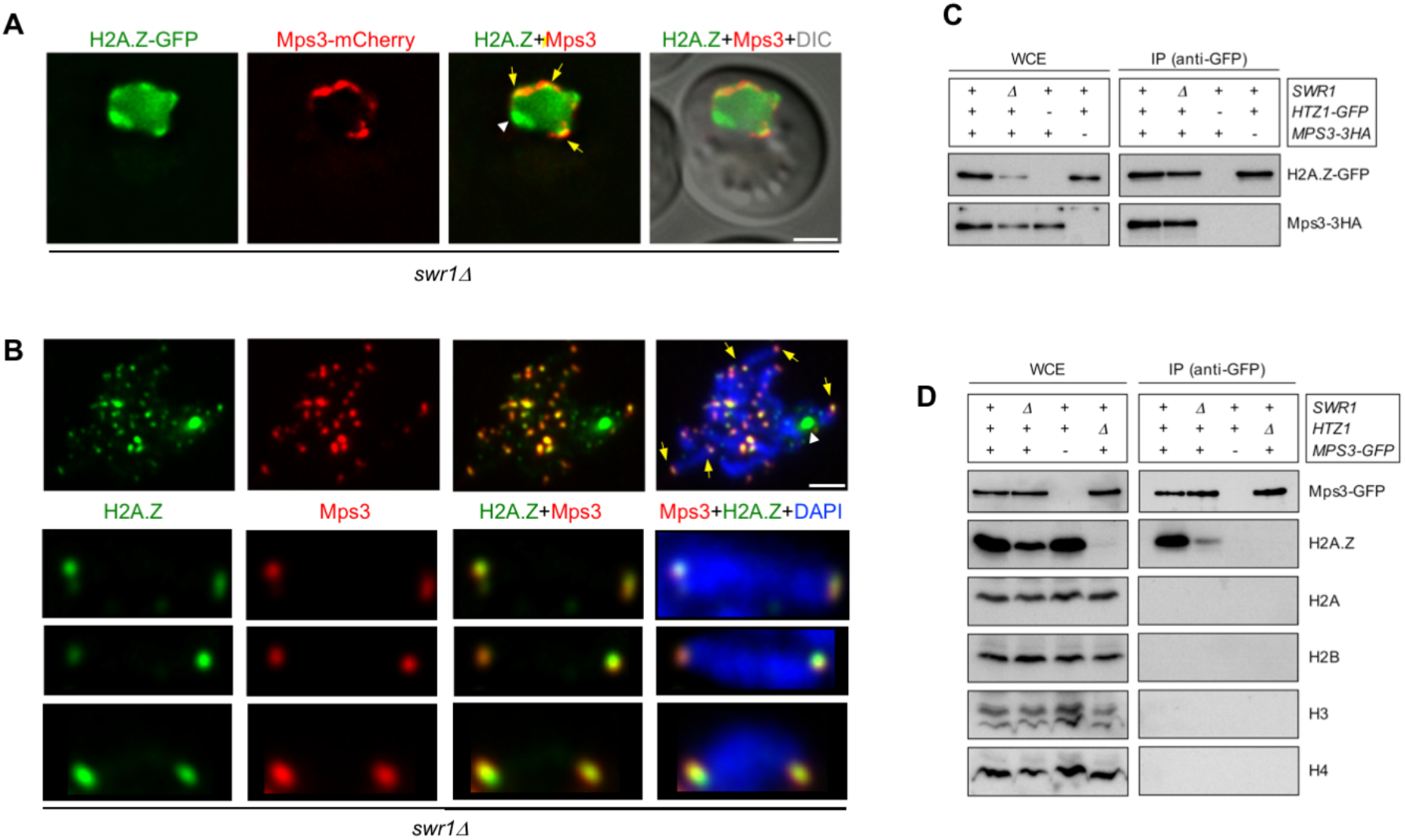
H2A.Z interacts and colocalizes with Mps3 at telomeres. **(A)** Images from a representative *swr1Δ* live cell expressing *HTZ1-GFP* and *MPS3-mCherry* 16 h after meiotic induction. Yellow arrows point to areas of the nuclear periphery where H2A.Z and Mps3 display colocalization. The white arrowhead points to the presumed accumulation of H2A.Z at the nucleolus. The differential interference contrast (DIC) image is shown as a reference for the cell outline. Scale bar, 2 μm. **(B)** Immunofluorescence of spread pachytene chromosomes from the *swr1Δ* mutant stained with DAPI to visualize chromatin (blue), anti-GFP to detect H2A.Z (green), and anti-mCherry to detect Mps3 (red). A representative nucleus is shown. The bottom panels display selected individual chromosomes. Spreads were prepared 16 h after meiotic induction. Yellow arrows point to some telomeric foci showing H2A.Z and Mps3 colocalization. The white arrowhead marks the accumulation of H2A.Z at the nucleolar rDNA region in *swr1Δ*. Scale bar, 2 μm. The strains used in (A, B) are DP1108 (*HTZ1-GFP MPS3-mCherry swr1Δ*) and DP1395 (*HTZ1-GFP MPS3-3HA swr1Δ*), respectively. **(C)** Whole cell extracts (WCE) prepared 16 h after meiotic induction were immunoprecipitated using GFP-Trap beads. WCEs and immunoprecipitates (IP) were analyzed by western blot using anti-GFP antibodies (to detect H2A.Z) and anti-HA antibodies (to detect Mps3). Strains used are: DP1394 (*SWR1 HTZ1-GFP MPS3-3HA*), DP1395 (*swr1Δ HTZ1-GFP MPS3-3HA*), DP1330 (*SWR1 HTZ1 MPS3-3HA*) and DP840 (*SWR1 HTZ1-GFP MPS3*). **(D)** WCEs prepared 16 h after meiotic induction were immunoprecipitated using GFP-Trap beads. WCEs and immunoprecipitates (IP) were analyzed by western blot using anti-GFP antibodies (to detect Mps3), and anti-H2A.Z, anti-H2A, anti-H2B, anti-H3 and anti-H4 histone antibodies. Strains used are: DP866 (*SWR1 HTZ1 MPS3-GFP*), DP1102 (*swr1Δ HTZ1 MPS3-GFP*), DP421 (*swr1 HTZ1 MPS3*) and DP867 (*SWR1 HTZ1ΔMPS3-GFP*).

### H2A.Z and Ndj1 interact at the nuclear periphery

We used the Bimolecular Fluorescence Complementation technique (BiFC) to further explore the physical interaction between H2A.Z and other LINC-associated components, such as Ndj1. BiFC permits direct visualization of protein interaction in living cells based on the association between two nonfluorescent fragments of a fluorescent protein brought in proximity by the interaction between proteins fused to the fragments (Kerppola, 2008; Miller et al., 2015). We found that H2A.Z and Ndj1 interact at the NE, as manifested by the reconstitution of fluorescence from the Venus variant of the Yellow Fluorescent Protein (Venus^YFP^) at the nuclear periphery in meiotic cells simultaneously expressing both moieties of Venus^YFP^ fused to H2A.Z and Ndj1 (*HTZ1-VN* and *NDJ1-VC*, respectively) (Figure 3A, 3D). Importantly, the H2A.Z-Ndj1 interaction was detected not only in the *swr1Δ* mutant, but also in the wild type (Figure 3B, 3D). This result indicates that the telomeric localization of H2A.Z is not an exclusive feature of the *swr1Δ* mutant, and also occurs in the wild type where it is masked in our cytological analysis of spread nuclei by the massive deposition of H2A.Z throughout the genome. We detected the reconstituted Venus^YFP^ signal in a fraction of cells in the culture (24.2% and 27.3% for wild type and *swr1Δ*, respectively) that roughly represents the population of prophase cells in the asynchronous BR strain background at the time point analyzed. In fact, a parallel meiotic culture used as a control for staging displayed ≈34% cells with Zip1-GFP signal at the same time point. Indeed, the use of a *ndt80Δ* mutation that prevents exit from prophase I increased the proportion of cells displaying H2A.Z-Ndj1 interaction to 54% and 55% in *SWR1* and *swr1Δ*, respectively, and the fraction of cells containing the Hop1-GFP prophase I marker to 70% (Figure S3). That is, the interaction was detected in ~80% of prophase cells. This is consistent with the observation that Ndj1-mediated tethering of telomeres to the NE only occurs in meiotic prophase I (Conrad et al., 2007). Of note, when the BiFC assay was performed in cells also expressing *MPS3-mCherry*, we observed that the Venus^YFP^ signal resulting from H2A.Z-Ndj1 interaction largely colocalized with Mps3 foci, thus confirming that it occurs at telomere attachment sites at the NE (Figure 3C; upper row).

**Figure 3.**
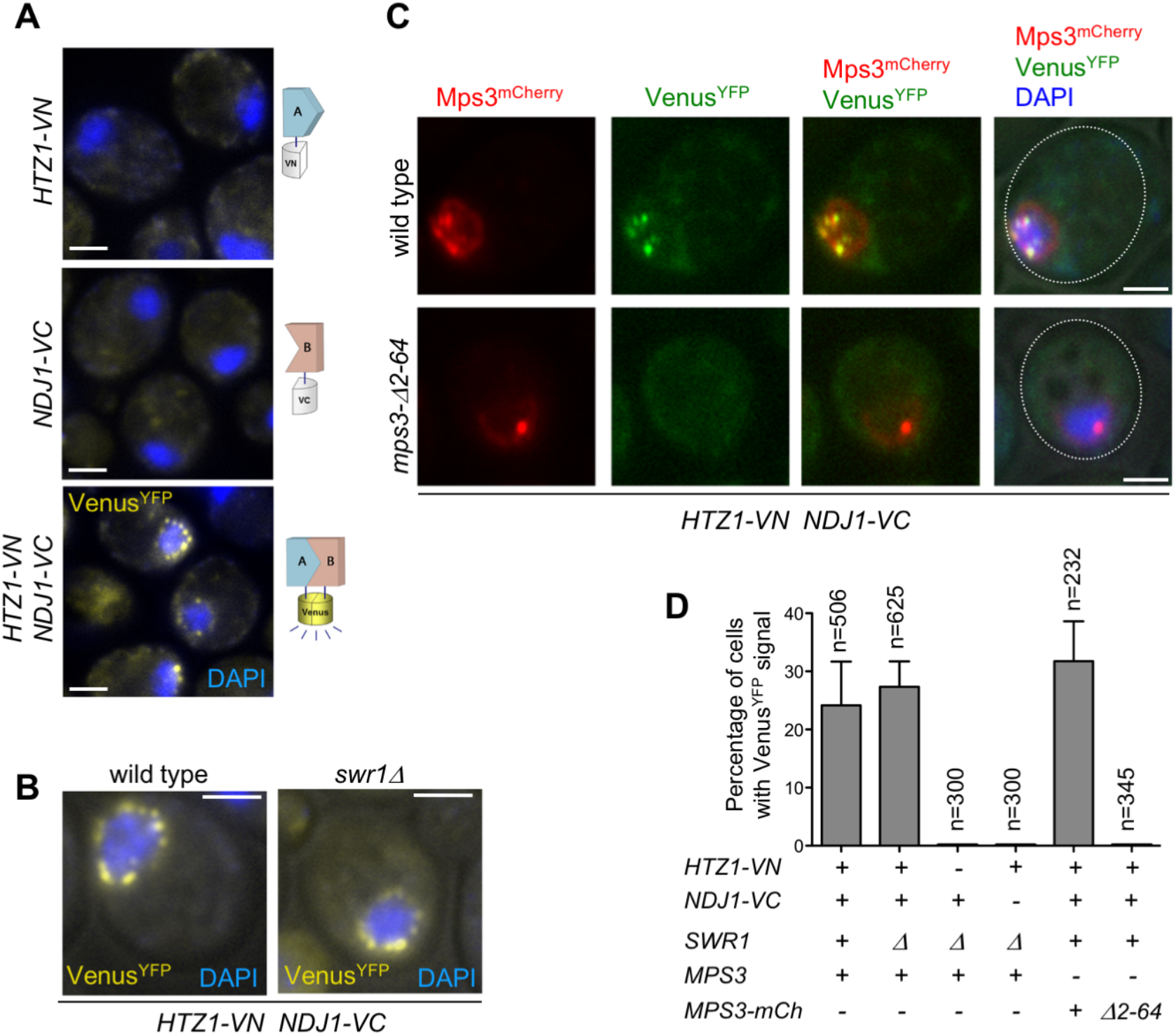
Analysis of H2A.Z and Ndj1 interaction by Bimolecular Fluorescence Complementation (BiFC) assay. **(A-B)** H2A.Z and Ndj1 interact at the nuclear periphery. Microscopy fluorescence images of cells expressing *HTZ1* fused to the N-terminal half of the Venus^YFP^ (*VN*) and/or *NDJ1* fused to the C-terminal half of the Venus^YFP^ (*VC*) as indicated. Nuclei are stained with DAPI (blue). The reconstitution of Venus^YFP^ fluorescence resulting from H2A.Z-VN/Ndj1-VC interaction appears in yellow. Images were taken 16 h after meiotic induction. Strains in (A) are: DP1540 (*htz1-VN*), DP1541 (*NDJ1-VC*) and DP1493 (*htz1-VN NDJ1-VC swr1Δ*). Strains in (B) are DP1496 (*htz1-VN NDJ1-VC*) and DP1493 (*htz1-VN NDJ1-VC swr1Δ*). **(C)** H2A.Z and Ndj1 interaction at the nuclear periphery depends on the 2-64 N-terminal domain of Mps3. Mps3-mCherry signal is shown in red, Venus^YFP^ in green, and DAPI in blue. Cells were imaged 16 h after meiotic induction. Strains in (C) are: DP1511 (*MPS3-mCherry htz1-VN NDJ1-VC*) and DP1512 (*mps3-Δ2-64-mCherry htz1-VN NDJ1-VC*). A single medial plane is shown in (A-C). **(D)** Quantification of the percentage of cells displaying Venus^YFP^ fluorescent signal in the experiments shown in (A-C). The analysis was performed in triplicate. The total number of cells scored (n) is shown. Error bars, SD. Scale bar, 2 μm.

### Telomeric localization of H2A.Z depends on LINC functional integrity

We next examined H2A.Z localization in mutants that compromise LINC-dependent telomere attachment to the NE, such as *mps3-Δ2-64* and *ndj1Δ*. The *mps3-Δ2-64* mutant lacks the 2-64 amino acids of Mps3 N-terminal domain. In the wild-type Mps3 protein this region protrudes into the nuclear inside serving as telomeric docking site via the Ndj1 protein (Conrad et al., 2007); therefore, in both *mps3-Δ2-64* and *ndj1Δ* mutants, telomere anchoring to the NE is impaired (Figure 4, left panels). We carried out this analysis in a *swr1Δ* mutant to get rid of the massive deposition of H2A.Z throughout chromatin enabling us to detect its telomeric localization. We found that the localization of H2A.Z at telomeres was lost in *swr1Δ mps3-Δ2-64* and *swr1Δ ndj1Δ* spread pachytene nuclei (Figure 4). Moreover, the H2A.Z-Ndj1 interaction detected by BiFC was abolished in the *mps3Δ2-64* mutant (Figure 3C, 3D), further supporting the notion that H2A.Z is recruited to an intact LINC complex. Of note, the presence of H2A.Z in the nucleolar vicinity observed in *swr1Δ* was maintained in the *swr1Δ mps3-Δ2-64* and *swr1Δ ndj1Δ* double mutants (Figure 4, arrowheads). Likewise, a strong H2A.Z focus not associated with the chromosomes that likely corresponds to the SPB (see below) was also detected (Figure 4, yellow arrows). Thus, these observations suggest that the association of H2A.Z to the telomeric regions specifically requires functional anchoring of the chromosomes to the NE mediated by the inner components of LINC.

**Figure 4.**
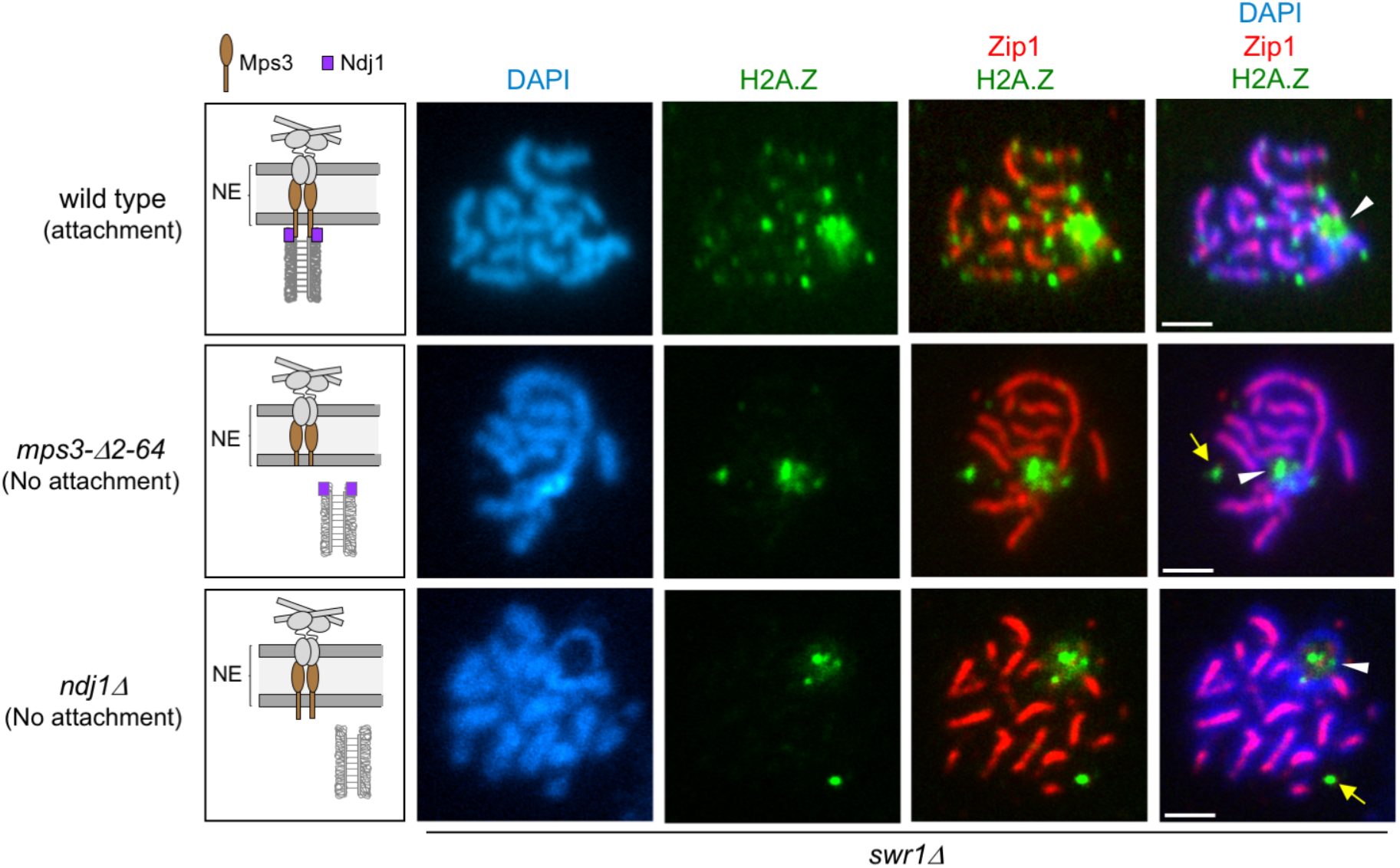
H2A.Z localization at chromosome ends requires telomere attachment to the NE. Immunofluorescence of spread pachytene nuclei from the *swr1Δ* mutant stained with DAPI to visualize chromatin (blue), anti-GFP to detect H2A.Z (green), and anti-Zip1 to mark the SC central region (red). White arrowheads mark the rDNA region lacking Zip1. Yellow arrows point to H2A.Z foci likely corresponding to the SPB (see text). The cartoons at the left schematize the LINC complex and the status of telomeric attachment in the different situations analyzed. Scale bar, 2 μm. Strains are DP1182 (wild type), DP1280 (*mps3-Δ2-64*) and DP1305 (*ndj1Δ*). 26, 28 and 24 nuclei were examined for wild type, *mps3-Δ2-64* and *ndj1Δ*, respectively.

### H2A.Z also colocalizes with Ndj1 and Mps3 at the SPB during meiosis

Several studies have shown that, in addition to telomeres, Mps3 and Ndj1 are also localized at the SPB in meiotic cells (Li et al., 2015; Rao et al., 2011). Since we have observed colocalization and interaction between H2A.Z and Mps3/Ndj1 at telomeres, we examined whether H2A.Z is also targeted to the SPB. In *swr1Δ* live meiotic cells, we observed that one of the peripheral spots of H2A.Z-GFP colocalized with the SPB core component Cnm67-mCherry (Figure 5A). Moreover, BiFC analysis revealed that one of the DAPI-surrounding foci where H2A.Z and Ndj1 interact corresponds to the SPB, as shown by colocalization of the Venus^YFP^ signal with Spc110, another SPB component, tagged with RedStar2 (Figure 5B). Immunofluorescence of spread meiotic chromosomes also showed colocalization between Cnm67 and H2A.Z at a defined focus (Figure 5C). However, in contrast to the telomeric localization of H2A.Z (Figure 4), the presence of H2A.Z at the SPB was maintained in *mps3Δ2-64* and *ndj1Δ* mutants during meiotic prophase, as manifested by the detection of a single H2A.Z focus associated to the characteristic monopolar prophase I spindle stained with tubulin antibodies (Figure S4). Thus, like in telomeres, our results suggest that Mps3, Ndj1 and H2A.Z also interact at the SPB, but our findings reflect differential requirements for targeting H2A.Z to the various subcellular locations.

**Figure 5.**
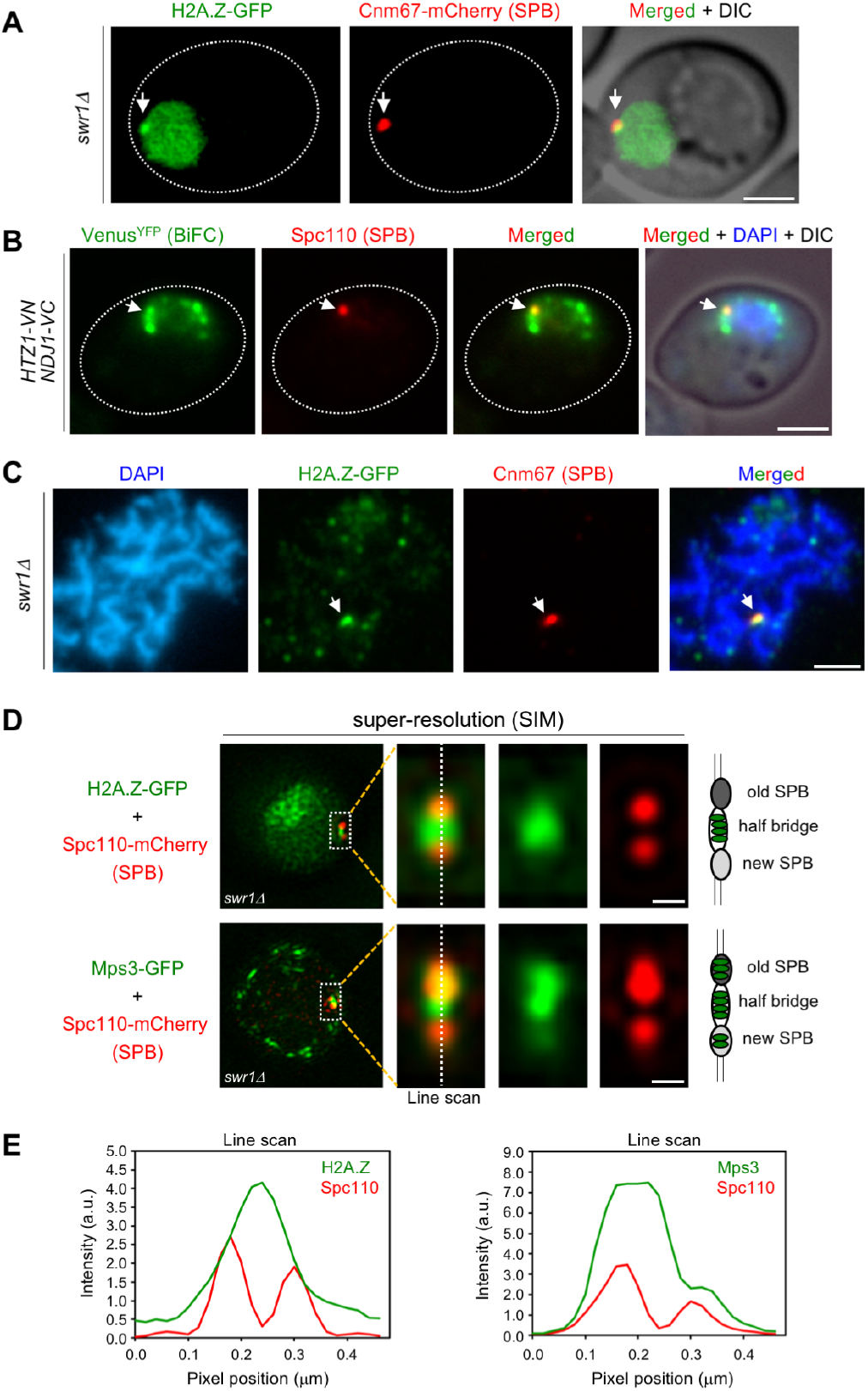
H2A.Z localizes to the SPB half bridge during meiotic prophase I. **(A)** Microscopy fluorescence image of a representative *swr1Δ* cell displaying a peripheral concentrated focus (arrow) of H2A.Z-GFP (green) colocalizing with the SPB marker Cnm67-mCherry (red). Images were taken from 16 h meiotic cultures. Scale bar, 2 μm. The strain is DP1172. **(B)** BiFC analysis of Venus^YFP^ fluorescence (green) reconstituted from H2A.Z-VN/Ndj1-VC interaction in cells also expressing the SPB marker Spc110-RedStar2 (red). Nuclei were stained with DAPI (blue). A representative cell is shown. The arrow points to a single BiFC Venus^YFP^ focus colocalizing with the SPB. Scale bar, 2 μm. The strain is DP1506. **(C)** Immunofluorescence of a spread pachytene representative nucleus stained with DAPI to visualize chromatin (blue), anti-GFP to detect H2A.Z (green), and anti-mCherry to mark the SPB (red). The arrow points to an H2A.Z focus colocalizing with Cnm67 (SPB). Scale bar, 2 μm. The strain is DP1172. 25 nuclei were examined. **(D)** Structured-Ilumination Microscopy (SIM) fluorescence images of representative *swr1Δ* cells expressing Spc110-mCherry (red) and H2A.Z-GFP (top images) or Mps3-GFP (bottom images), in green. Scale bar, 0.1 μm. **(E)** Average intensity of the indicated proteins along the depicted line scan in all cells analyzed in (D). Strains are DP1578 (*HTZ1-GFP SPC110-mCherry*) and DP1576 (*MPS3-GFP SPC110-mCherry*); 33 and 26 cells were examined, respectively.

### H2A.Z localizes to the SPB half bridge

To determine the precise localization of H2A.Z within the SPB structure during meiotic prophase, we used Structured Illumination Microscopy (SIM). We examined the colocalization of H2A.Z-GFP, as well as Mps3-GFP for comparison, with the Spc110-mCherry protein, a component of the SPB inner plaque (Figure 5D). We focused on prophase cells containing duplicated unseparated SPBs. The old and new SPBs could be distinguished by the stronger and weaker Spc110-mCherry signal, respectively (Burns et al., 2015). Most of the H2A.Z-GFP signal concentrated in the area in between both SPBs that corresponds to the half bridge, and only a limited overlap with the SPBs was observed (Figure 5E, left graph). Like H2A.Z, Mps3-GFP was also detected in the half bridge, but also displayed more extensive colocalization with Spc110-mCherry (Figure 5E, right graph), consistent with the idea that Mps3 is a dual component of the bridge and the membrane domain that surrounds the SPB core (Chen et al. 2019). Thus, we conclude that the fraction of H2A.Z present in the SPB mainly localizes to the half bridge structure that tethers duplicated SPBs during meiotic prophase I.

### Altered distribution of Mps3 along the NE in the absence of H2A.Z

To further explore the meiotic relationship between Mps3 and H2A.Z, we examined Mps3 levels and localization in the *htz1Δ* mutant. Western blot analysis of Mps3 production in meiotic cultures showed that the protein was heavily induced during meiotic prophase and then declined at late time points as meiosis and sporulation progresses (Figure 6A). The dynamics of Mps3 production was similar in wild type and *htz1Δ*, but global Mps3 levels were reduced in the *htz1Δ* mutant. To rule out the possibility that the reduction in the amount of Mps3 was exclusively due to an inefficient meiotic progression in *htz1Δ* (Gonzalez-Arranz et al., 2018), we measured Mps3 levels in the prophase-arrested *ndt8OΔ* mutant, monitoring also Mek1 production as a proxy for a meiotic prophase I protein (Ontoso et al., 2013). This analysis revealed that the lack of H2A.Z specifically affects Mps3, but not Mek1, global levels (Figure S5). By immunofluorescence of pachytene chromosome spreads, we found that Mps3 remained at telomeres in the absence of H2A.Z. However, Mps3 telomeric localization was lost in the *ndj1Δ* mutant used as control for comparison (Conrad et al., 2007) (Figure 6B). Thus, although H2A.Z telomeric localization depends on the N-terminal domain of Mps3 (see above), the anchoring of Mps3 to telomeres is independent of H2A.Z. We also examined Mps3-GFP localization in live meiotic cells. The presence of Hop1-mCherry signal was used to stage cells in prophase I (Figure 6C). Analysis of the pattern of Mps3-GFP localization in whole meiotic prophase cells revealed foci with a rather uniform distribution along the NE in the wild type. In contrast, Mps3 showed a more irregular distribution and it appeared to be more concentrated at defined NE regions at a higher frequency in the *htz1Δ* mutant (Figure 6C, 6D; Video S1). We conclude that H2A.Z is required for proper distribution of Mps3 along the NE during meiotic prophase I, but not for telomere attachment.

**Figure 6.**
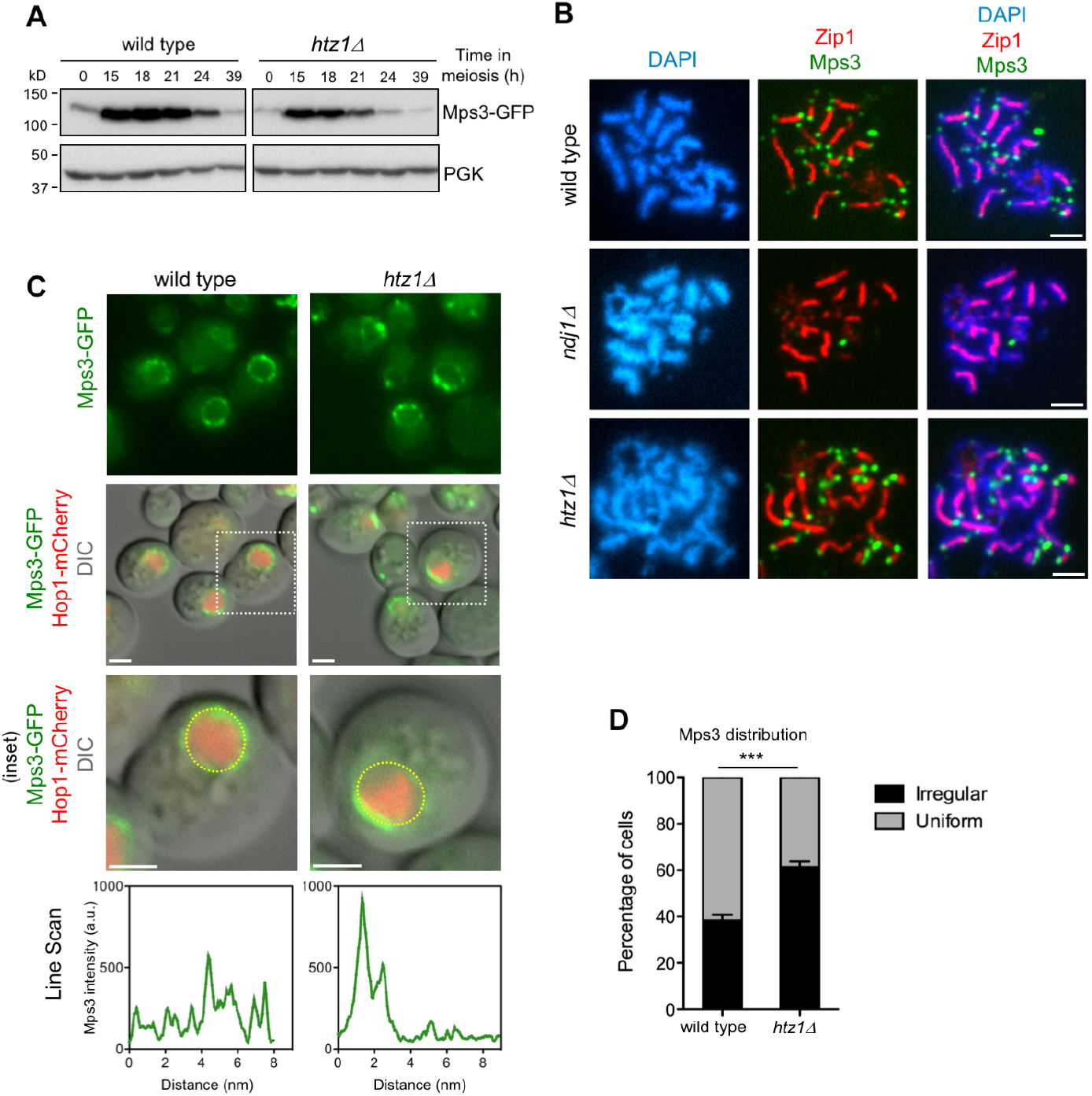
Altered levels and distribution of Mps3 in the absence of H2A.Z. **(A)** Western blot analysis of Mps3 production during meiosis detected with anti-GFP antibodies. PGK was used as a loading control. Strains in (A) are: DP866 (wild type) and DP867 (*htz1Δ*). **(B)** Immunofluorescence of spread pachytene nuclei stained with DAPI to visualize chromatin (blue), anti-GFP to detect Mps3 (green), and anti-Zip1 to mark the SC central region (red). Scale bar, 2 μm. Strains in (B) are: DP866 (wild type), DP1103 (*ndj1Δ*) and DP867 (*htz1Δ*). **(C)** Microscopy fluorescence images of cells expressing *MPS3-GFP* and *HOP1-mCherry*. The presence of Hop1-mCherry was used to detect meiotic prophase cells. Stacks of images in the Z-axis were taken, but a single central plane from representative cells is shown. The line scan plots represent the GFP signal along the depicted yellow circle line in the bottom row cells. Scale bar, 2 μm. **(D)** The distribution of Mps3 was analyzed in maximum-intensity projections from images obtained as in (C). Two categories were established: uniform and irregular. Cells scored as “uniform” display Mps3-GFP signal homogeneously distributed. Cells scored as “irregular” display Mps3-GFP signal concentrated to one area of the NE. Only cells displaying Hop1-mCherry signal were considered in the analysis. This quantification was performed in triplicate. A total of 383 and 387 cells were scored for wild type and *htz1Δ*, respectively. Strains in (C-E) are: DP1032 (wild type) and DP1033 (*htz1Δ*).

### Chromosome motion is reduced in the absence of H2A.Z

One of the main meiotic functions of the LINC complex is to promote telomere-led chromosome movement during prophase I (Conrad et al., 2008; Koszul et al., 2008). Since we found that H2A.Z interacts with LINC components during meiosis we hypothesized that H2A.Z could also contribute to meiotic chromosome motion. Initially, we used strains expressing *ZIPl-GFP* to follow chromosome movement, as previously described (Scherthan et al., 2007; Sonntag Brown et al., 2011). In addition to wild-type and *htz1Δ* strains, we also analyzed the *ndj1Δ* mutant as a control for defective chromosome mobility and the *swr1Δ* mutant in which H2A.Z is not deposited into the chromatin (see above). To minimize experimental variation, we mixed wild-type and mutant cells from meiotic cultures (16 h) in the same microscopy culture chamber to analyze chromosome movement in parallel. Wild-type cells could be easily distinguished by the presence of Pma1-mCherry, a plasma membrane protein that was tagged to mark these cells (Figure 7A). We tracked the ends of individual synapsed chromosomes and measured the distance traveled during a defined time (Figure 7B; Video S2). We found that the average velocity of chromosome movement was reduced in the *htz1Δ* mutant, although to a lesser extent than in *ndj1Δ*. In contrast, the *swr1Δ* mutant was not affected (Figure 7C).

**Figure 7.**
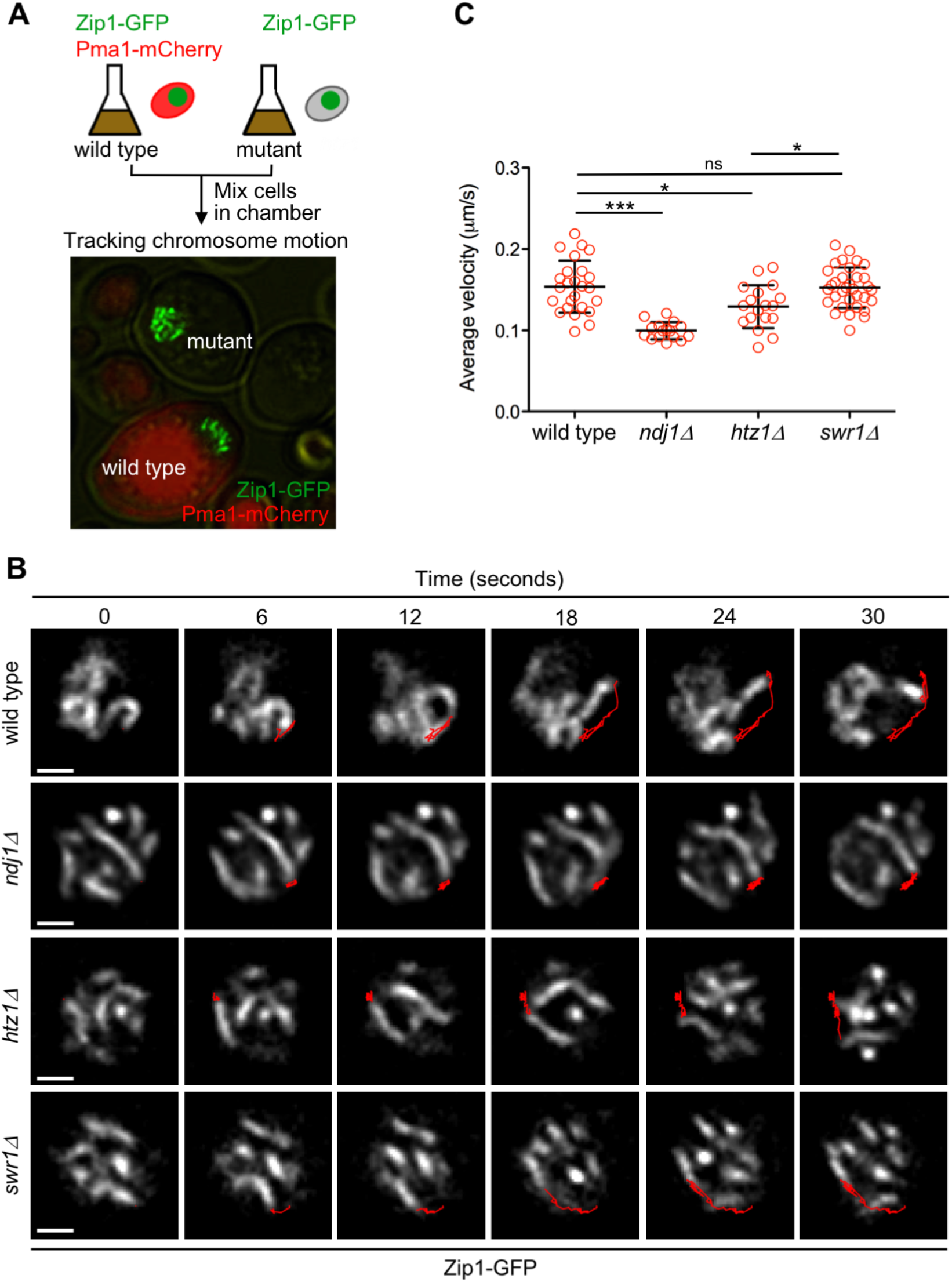
H2A.Z drives prophase chromosome movement. **(A)** Experimental setup to monitor movement of Zip1-GFP labeled chromosomes. Aliquots of cells from 16-hour meiotic cultures of wild type and *ndj1Δ, htz1Δ* or *swr1Δ*, mutants were mixed in microscopy chambers and followed in parallel by time-lapse fluorescence microscopy. Wild-type cells were distinguished by the presence of Pma1-mCherry. **(B)** Representative images of nuclei from the indicated strains at different time intervals. Zip1-GFP signal is shown. The red line represents the path traveled by an individual chromosome end throughout the time lapse. Scale bar, 1 μm. **(C)** Quantification of the average velocity of chromosome movement. Mean and SD are represented. Strains are DP1057 (wild type), DP957 (*ndj1Δ*), DP838 (*htz1Δ*) and DP1091 (*swr1Δ*). 25, 16, 18 and 33 chromosome measurements from different cells of wild type, *ndj1Δ, htz1Δ* and *swr1Δ*, respectively, were performed in three independent time-lapse experiments.

Velocity measurements based on Zip1-GFP rely on the ability to track a single chromosome pair in the maze of all synapsed chromosomes, which is not always possible. Therefore, for a more extensive and accurate analysis of telomere-driven movement we used a *tetO-tetR* system (Clemente-Blanco et al., 2011) to generate strains harboring the left telomere of chromosome IV (*TEL4L*) labeled with GFP (Figure 8A). These strains also expressed *ZIP1-mCherry* as a marker for prophase I stage and synapsed chromosomes (Figure 8B, 8C). In addition, we also introduced the *P_CUP1_-IME1* construct to increase the synchrony of the meiotic cultures (Chia and van Werven, 2016). *TEL4L* trajectory was tracked in time-lapse experiments of wild-type, *ndj1Δ, htz1Δ* and *swr1Δ* strains (Figure 8D; Video S3). Measurement of both average and maximum velocity of *TEL4L* movement during prophase I using this system also revealed that chromosome motion was significantly reduced in the *htz1Δ* mutant (Figure 8E, 8F; Figure S6). Interestingly, consistent with the chromatin-independent interaction between H2A.Z and the LINC complex, the *swr1Δ* mutant did not display reduced mobility. As expected, the *ndj1Δ* mutant showed a dramatic reduction in *TEL4L* movement (Figure 8E, 8F; Figure S6). We conclude that H2A.Z is a novel LINC-associated component required for proper chromosome motion during meiotic prophase I.

**Figure 8.**
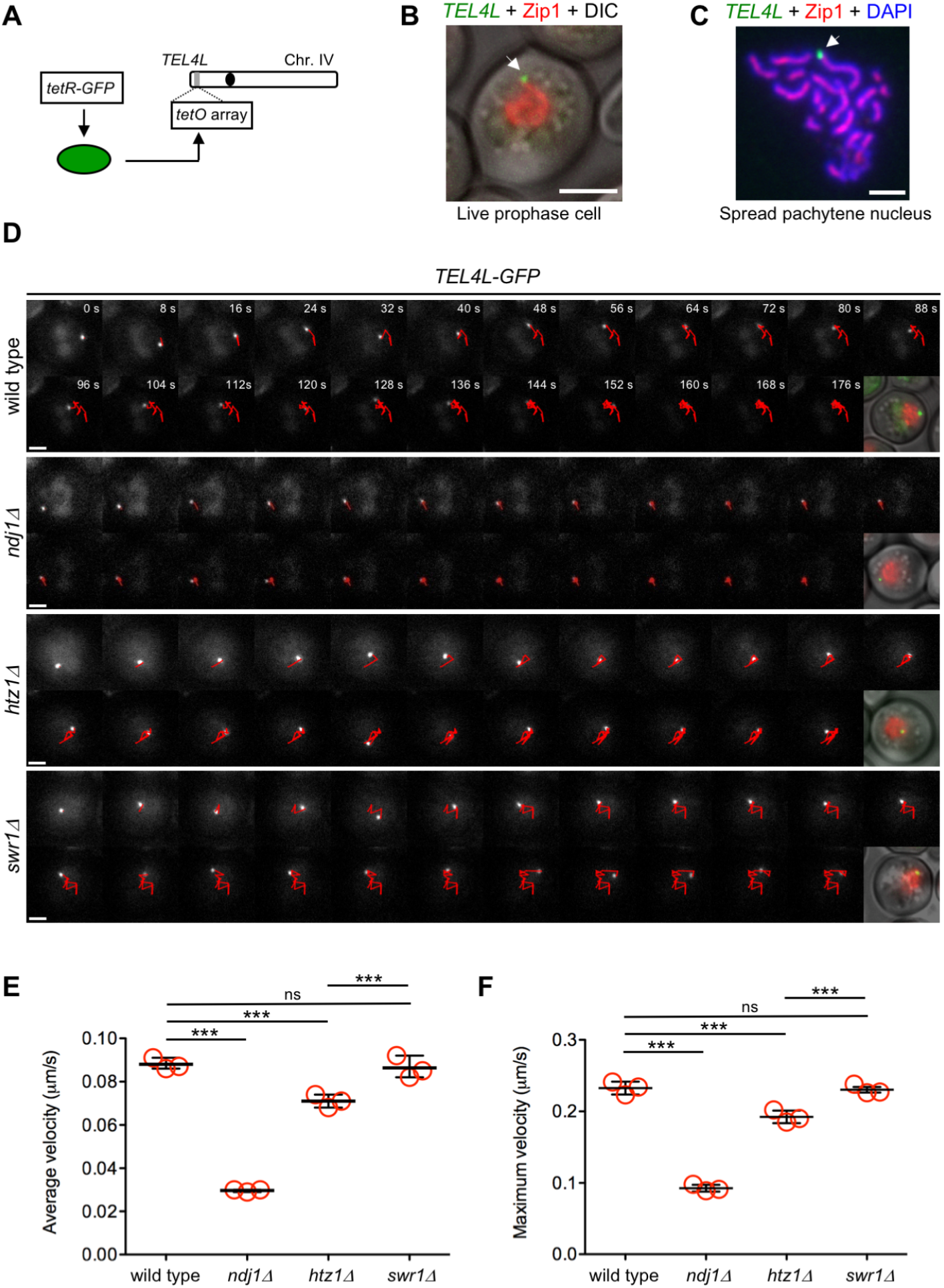
H2A.Z, but not SWR1, is required for proper telomere motion during meiotic prophase. **(A)** Schematic representation of GFP tagging of the left telomere of chromosome IV (*TEL4L*). **(B)** Representative image of a live prophase cell expressing *ZIP1-mCherry* (red) with *TEL4L* (arrow) labeled with GFP (green). Scale bar, 2 μm. **(C)** Representative image of a spread pachytene nucleus stained with anti-Zip1 (red), anti-GFP (*TEL4L*; green) and DAPI (blue). Arrow points to *TEL4L*. Scale bar, 2 μm. **(D)** Representative time-lapse fluorescence images of *TEL4L-GFP* at different time intervals (expressed in seconds at the top panels). The red line depicts the path traveled by *TEL4L-GFP* throughout the time lapse. A merge image of DIC, Zip1-mCherry (red) and *TEL4L-GFP* (green) is shown after the last frame. Scale bar, 2 μm. **(E)** Quantification of average velocity of *TEL4L* movement. **(F)** Quantification of maximum velocity of *TEL4L* movement. In (E) and (F), the mean values from three independent experiments are plotted in the graphs. Data from each individual experiment and all multiple statistical comparisons are presented in Figure S6. A total of 101, 146, 172 and 139 measurements of *TEL4L* velocity from wild type, *ndj1Δ, htz1Δ* and *swr1Δ*, respectively, were performed in the three independent time-lapse experiments. Strains used in (D-F) are DP1692 (wild type), DP1722 (*ndj1Δ*), DP1693 (*htz1Δ*) and DP1694 (*swr1Δ*).

## DISCUSSION

The H2A.Z histone variant is involved in a myriad of biological processes, both in mitotic and meiotic cells, that rely on its chromatin incorporation at particular genomic positions where the ATP-dependent chromatin remodeler SWR1 is responsible for replacing H2A-H2B dimers to H2A.Z-H2B at nucleosomes (Billon and Cote, 2013). In this work, we characterize in detail an alternative localization of H2A.Z in different sub-compartments of meiotic cells that is independent of SWR1 and, hence, of chromatin. Indeed, our cytological studies of H2A.Z in the *swr1Δ* mutant allowed us to uncover additional locations of H2A.Z (chromosome ends and SPB) that are otherwise masked in wild-type cells due to the widespread incorporation of H2A.Z throughout chromatin. Like in mitotic cells (Raisner et al., 2005), ChIP-seq analysis of H2A.Z distribution during meiotic prophase has confirmed the absence of H2A.Z chromatin deposition throughout the genome during meiosis in the *swr1Δ* mutant; in particular, at gene promoter regions. Here, we describe chromosome movement as a novel SWR1-independent meiotic function for this histone variant.

Prompted by our cytological analysis of H2A.Z in the *swr1Δ* mutant during meiotic prophase revealing a telomeric localization and by an earlier report describing the interaction between H2A.Z and Mps3 in vegetative cells (Gardner et al., 2011), we explored the relationship of H2A.Z with LINC components and LINC-associated components, such as the SUN-domain protein Mps3 and the meiosis-specific telomeric protein Ndj1, respectively. The colocalization of H2A.Z with Mps3 at meiotic telomeres and the physical interaction between H2A.Z and both Mps3 and Ndj1, particularly at the NE, strongly suggests that H2A.Z is an additional LINC-associated factor (Figure 1A). It has been proposed that H2A.Z may contribute to the nuclear trafficking of Mps3 in mitotic cells (Gardner et al., 2011); however, we found that Mps3 is still detectable at chromosome ends in the *htz1Δ* mutant, indicating that Mps3 does not require H2A.Z to be delivered to the NE in meiotic cells. Since Mps3 is embedded in the NE, its detection on nuclear spread preparations depends on the attachment of the N-terminal domain to the telomeres via Ndj1 (Conrad et al., 2007). The fact that Mps3 telomeric localization is maintained in the *htz1Δ* mutant also indicates that telomere attachment is not disrupted in the absence of H2A.Z. Nevertheless, we found that the distribution of Mps3 throughout the NE is altered in the *htz1Δ* mutant that often displays an aberrant confinement of Mps3 towards one side of the nucleus. Thus, unlike mitotic cells, Mps3 does not require H2A.Z to reach the NE during meiosis, but H2A.Z is required to sustain homogenous distribution of Mps3 along the NE. This accumulation of Mps3 observed in *htz1Δ* is reminiscent of the transient meiotic bouquet (Trelles-Sticken et al., 1999), suggesting that H2A.Z may facilitate the dispersion of telomeres after the bouquet-like stage. The telomeric colocalization and the physical interaction between H2A.Z and Mps3/Ndj1 strongly suggest that H2A.Z directly impinges on LINC dynamics.

We found that disruption of telomere attachment, either by deleting *NDJ1* or eliminating the 2-64 amino acids of the Mps3 N-terminal domain, prevents localization of H2A.Z to chromosome ends. This observation suggests that although H2A.Z is not necessary to anchor the telomeres to the NE, it may reinforce the tethering to support robust chromosome movement. A similar scenario has been described in the *mps3-dCC* mutant lacking a fragment of an internal domain of Mps3 located in the lumen of the NE. Like in *htz1Δ*, telomeric attachment is maintained in *mps3-dCC*, but chromosome movement is mildly affected (Lee et al., 2012). Consistent with this notion, the reduction in chromosome velocity detected in *htz1Δ* is not as dramatic as in the *ndj1Δ* mutant where telomere attachment via Mps3 is lost (Conrad et al., 2007). Incorporation of H2A.Z into nucleosomes produces changes in chromatin rigidity (Gerhold et al., 2015; Neumann et al., 2012); thus, it is formally possible that the reduced mobility of *htz1Δ* chromosomes could stem from altered chromatin compaction preventing proper transmission of the forces generated by the actin cytoskeleton to the chromosomes. However, the *swr1Δ* mutant shows no defect in chromosome movement indicating that chromatin deposition of H2A.Z has little impact on this phenomenon. In fact, the interaction between Mps3 and H2A.Z, and between Ndj1 and H2A.Z, persists in the absence of SWR1 both in mitotic (Gardner et al., 2011) and meiotic cells (this work). We hypothesize that the irregular accumulation of Mps3 at certain areas of the NE detected in *htz1Δ* cells may interfere with proper telomere-led movement. The involvement of H2A.Z in chromatin movement has been reported also in mitotic cells where H2A.Z promotes the recruitment of unrepairable DSBs to the NE by Mps3 anchoring (Horigome et al., 2014; Kalocsay et al., 2009). However, this nuclear relocalization relies on SWR1-dependent chromatin deposition of H2A.Z, suggesting that different mechanisms are involved.

Certain histone post-translational modifications, such as Set1-mediated H3K4 methylation have been shown to be involved in bouquet formation and telomere redistribution throughout the NE (Trelles-Sticken et al., 2005). Like H2A.Z, Set1 is also involved in the regulation of the so-called telomere-position effect (TPE) (Krogan et al., 2002; Martins-Taylor et al., 2011; Meneghini et al., 2003). However, the contribution of Set1 and H2A.Z to meiotic nuclear dynamics likely derives from different mechanisms. First, we show here that, unlike TPE (Kobor et al., 2004; Krogan et al., 2003; Mizuguchi et al., 2004), the interaction of H2A.Z with the LINC complex and its role in promoting chromosome movement does not require its SWR1-dependent chromatin incorporation. Second, the telomere dispersion defect of *set1Δ* mutants appears to be independent of Ndj1 (Trelles-Sticken et al., 2005).

In addition to the telomeric localization, we also describe here the presence of H2A.Z in another cellular structure devoid of chromatin, the SPB, in particular, the half bridge where Mps3 is also located. However, the targeting of H2A.Z to the SPB presents different requirements because, unlike its telomeric localization, it does not require Ndj1 or the 2-64 amino acids of the Mps3 N-terminal domain. Notably, we detect interaction between H2A.Z and Ndj1 in the wild type (Figure 5B), but not in the *mps3Δ2-64* mutant (Figure 3C) consistent with the observation that Ndj1 requires the N-terminal domain of Mps3 for SPB recruitment (Li et al., 2015). What could be the function of H2A.Z at the SPB? During meiotic prophase I, Ndj1, which is recruited to the SPB by Mps3, protects the cohesion between duplicated SPBs (Li et al., 2015). Phosphorylation of Mps3 at S70 promotes the proteolytic cleavage of the protein enabling irreversible separation of sister SPBs (Li et al., 2017). We speculate that the presence of H2A.Z at the SPB at the same time and location as Mps3 and Ndj1 may be indicative of a role for H2A.Z in SPB dynamics; future experiments will address this question. The fact that Mps3, Ndj1 and, as described here, also H2A.Z, colocalize and interact both at telomeres and SPB is consistent with a number of observations indicating that proper nuclear architecture and LINC-mediated contacts between chromosomes and the NE are important to coordinate interhomolog interactions with subsequent chromosome segregation. In *S. pombe*, multiple lines of evidence support this coordination (Fennell et al., 2015; Fernandez-Alvarez et al., 2016; Katsumata et al., 2016; Tomita and Cooper, 2007).

In sum, we describe here a novel role for H2A.Z in telomere-led meiotic chromosome motion that is independent of its deposition on chromatin by SWR1. The budding yeast *htz1Δ* mutant displays various meiotic phenotypes including slower meiotic progression and reduced viability of meiotic products. Interestingly, although the *swr1Δ* mutant also shows meiotic defects, these phenotypes are less severe in *swr1Δ* compared to *htz1Δ*. Indeed, spore viability is 95% in the wild type, 76% in *htz1Δ* and 88% in *swr1Δ* (Gonzalez-Arranz et al., 2018), consistent with the notion that H2A.Z possesses additional meiotic roles unrelated to SWR1, namely chromosome motion. Thus, we propose that, at least in budding yeast, H2A.Z performs both chromatin-dependent and chromatin-independent functions all of them contributing to sustain accurate gametogenesis. Cytological analyses of H2A.Z localization in mice spermatocytes have revealed a dynamic spatiotemporal localization of this histone variant on different euchromatin and heterochromatin domains (sex body) suggestive of a functional impact on mammalian meiosis (Greaves et al., 2006; Ontoso et al., 2014). In the future, it will be interesting to determine whether the chromatin-independent function of H2A.Z is also evolutionarily conserved.

## MATERIALS AND METHODS

### Yeast strains

Yeast strain genotypes are listed in Table S1. All the strains are isogenic to the BR1919 background (Rockmill and Roeder, 1990). The *swr1::natMX4, swr1::hphMX4, ndj1::natMX4, ndj1::kanMX6, htz1::natMX4, mps3::natMX4 and mps3::hphMX4* gene deletions were made using a PCR-based approach (Goldstein and McCusker, 1999; Longtine et al., 1998). The *htz1::URA3* deletion and the functional *HTZ1-GFP* construct were previously described (Gonzalez-Arranz et al., 2018). The *MPS3-GFP, MPS3-3HA, NET1-RedStar2, SPC110-RedStar2, SPC110-mCherry, CNM67-mCherry, HOP1-mCherry* and *PMA1-mCherry* gene tagging constructs were also made by PCR (Janke et al., 2004; Longtine et al., 1998; Sheff and Thorn, 2004). To generate *P_CUP1_-IME1* strains, the 1760 bp promoter region of *IME1*, including the *IRT1* lcRNA (Chia and van Werven, 2016), was replaced by the *CUP1* promoter amplified from pYM-N1 (Janke et al., 2004). Strains producing a version of Mps3 lacking amino acids 2-64 of the N-terminal domain were created as follows. One allele of the essential *MPS3* gene was deleted in a diploid strain. The heterozygous *MPS3/mps3-hphMX4* diploid was transformed with the *URA3-based* pSS326 centromeric plasmid harboring *mps3Δ2-64*. 5-Fluoroorotic acid (FOA)-resistant and hygromycin-resistant spores were selected and further checked for the presence of *mps3Δ2-64* expressed from pSS326 as the only source of this protein in the cells. As a control, the same procedure was followed using the pSS269 plasmid expressing wild-type *MPS3*. The *HTZ1-VN* and *NDJ1-VC* strains used in the Bimolecular Fluorescence Complementation (BiFC) assay were constructed using the plasmids pFA6a-VN-TRP1 and pFA6a-VC-kanMX6 containing the N-terminal (VN) or C-terminal fragment (VC) of the Venus variant of yellow fluorescent protein (Sung and Huh, 2007). Strains carrying Tel4L marked with GFP were generated as follows. First, the *tetR-GFP* construct was integrated at *leu2* by transforming with the *Afl*II-digested pSS329 plasmid. Second, the *tetO(50*) array was inserted into a region close to the left telomere of chromosome IV (Tel4L) by transforming with the pSS330 plasmid cut with *AflII*. Strains producing *ZIP1* tagged with mCherry at position 700 were constructed using the *delitto perfetto* approach (Stuckey and Storici, 2013). Basically, a fragment containing *mCherry* flanked by 60-nt *ZIP1* sequences upstream and downstream of the codon for amino acid 700 was obtained by PCR from plasmid pSS266. This fragment was transformed into a strain carrying the CORE cassette (*URA3-kanMX4*) inserted at the position corresponding to amino acid 700 of the *ZIP1* gene. FOA-resistant and G418-sensitive transformants were obtained and correct clones were selected.

All constructs and mutations were verified by PCR analysis and/or DNA sequencing. The sequences of all primers used in strain construction are available upon request. All strains were made by direct transformation of haploid parents or by genetic crosses always in an isogenic background. Diploids were made by mating the corresponding haploid parents and isolation of zygotes by micromanipulation.

### Plasmids

The plasmids used in this work are listed in Table S2. To generate pSS266, a PCR fragment containing 468 nt of the *MPS3* promoter, the *MPS3-mCherry* C-terminal fusion and the *ADH1* terminator was amplified from genomic DNA of a strain harboring the *MPS3-mCherry* construct and blunt-cloned into the pJET2.1 vector (ThermoFisher). A *BglII-BglII* fragment from pSS266 containing *MPS3-mCherry* was then cloned into *Bam*HI of pRS424 to generate pSS267. Then, a 3.5-kb *XhoI-NotI* fragment from pSS267 containing *MPS3-mCherry* was cloned into the same sites of the centromeric vector pRS316 to generate pSS269. A version of *MPS3-mCherry* lacking the sequences encoding amino acids 2-64 of the N-terminal domain (*mps3Δ2-64*) was made by site-directed mutagenesis of pSS269 using divergent oligonucleotides flanking the region to be deleted to generate plasmid pSS326. The pSS329 plasmid contains the *tet* repressor fused to *NLS-GFP* and expressed from the *URA3* promoter (Michaelis et al., 1997). *Eco*RI or *AflII* digestion of pSS329 targets TetR-GFP at *leu2*. The pSS330 plasmid harbors an approximately 5.4-kb array with 50 tandem repeats of the *tetO* operator inserted between the *HXT15* and *THI13* genes located close to *TEL4L* cloned into *BamHI-XbaI* of pRS406.

### Meiotic cultures and synchronous sporulation of BR strains

To induce meiosis and sporulation, BR strains were grown in 3.5 ml of 2X Synthetic Complete medium (2% glucose, 0.7% yeast nitrogen base without amino acids, 0.05% adenine, and complete supplement mixture from Formedium at twice the particular concentration indicated by the manufacturer) for 20–24 h, then transferred to 2.5 ml of YPDA (1% yeast extract, 2% peptone, 2% glucose, 0.02% adenine) and incubated to saturation for additional 8 hr. Cells were harvested, washed with 2% potassium acetate (KAc), resuspended into 2% KAc (10 ml), and incubated at 30°C with vigorous shaking to induce meiosis. Both YPDA and 2% KAc were supplemented with 20 mM adenine and 10 mM uracil. The culture volumes were scaled up when needed.

To increase synchrony in the meiotic cultures for Tel4L-GFP tracking, BR strains containing *P_CUP1_-IME1* were used. Culture conditions during pre-sporulation were similar as described above, except that YPDA contained 1% glucose. Cells were transferred to 2% KAc and, after 12 h, CuSO4 was added at a final concentration of 50 μM to induce *IME1* expression and drive meiotic entry. Cells were imaged 6 h after *IME1* induction, when approximately 73% of cells in the culture contained linear stretches of Zip1-mCherry.

### Western blotting and immunoprecipitation

Total cell extracts for western blot analysis in Figure 6A and Figure S5 were prepared by TCA precipitation from 5-ml aliquots of sporulation cultures as previously described (Acosta et al., 2011). The antibodies used are listed in Table S3. The ECL, ECL2 or SuperSignal West Femto reagents (ThermoFisher Scientific) were used for detection. The signal was captured on films and/or with a ChemiDoc XRS system (Bio-Rad).

For co-immunoprecipitation experiments, cells from 200 ml of meiotic cultures (16 h after meiotic induction) were harvested and washed with extraction buffer (20 mM HEPES-NaOH pH 7.5, 300 mM NaCl, 1mM EDTA, 5 mM EGTA, 50 mM NaF, 50 mM β-glycerophosphate, 1mM DTT, 1mM PMSF) containing one tablet of EDTA-free Complete Protease Inhibitors (Roche). The cell pellet was frozen in liquid nitrogen and ground with a freezer mill (6775 Freezer Mill). Ground cell powder was allowed to thaw on ice and then resuspended in 9 ml of lysis buffer (extraction buffer plus 0.5% Triton X-100). After homogenization with a homogenizer (ultra-turrax T10 basic, IKA) for 30 s, lysates were centrifuged at 3000 x g for 10 min at 4°C, and the resulting supernatant was used for immunoprecipitation saving 100 μl for input analysis. 50 μl of GFP-Trap magnetic agarose (Chromotek) were added to the remaining lysate to immunoprecipitate GFP-tagged proteins. After 3h incubation with rotation at 4°C, beads were washed five times with extraction buffer and the bound proteins were eluted by boiling in 2X Laemmli buffer. Samples from both input lysates and immunoprecipitates were analyzed by SDS-PAGE followed by western blotting.

### Chromatin immunoprecipitation and Illumina sequencing

At the 0 and 15 hour time points, 7 mL meiotic cultures (OD_600_ ~6-7) were harvested and fixed for 30 min with 1% formaldehyde. The crosslinking reaction was quenched by the addition of 125mM glycine. Chromatin immunoprecipitation was performed as described (Blitzblau et al., 2012). Samples were immunoprecipitated with 3 μL polyclonal rabbit anti-GFP serum per IP. Library quality was confirmed by Qubit HS assay kit and 2200 TapeStation. 51-bp single-end sequencing was accomplished on an Illumina HiSeq 2500 instrument.

### Processing Illumina data

Sequencing reads were mapped to a high-quality assembly of S288C (Yue et al., 2017) using Bowtie (Langmead et al., 2009). Reads with up to 2 mismatches across all 51bp were considered during mapping, and reads with more than one reportable alignment were mapped to the best position. Reads were also mapped to the SK1 genome with similar results. Reads were extended towards 3’ ends to a final length of 200 bp using MACS-2.1.0 (https://github.com/taoliu/MACS) (Zhang et al., 2008) All pileups were SPMR-normalized (signal per million reads) and fold-enrichment of the ChIP data over the input data was calculated. The 95% confidence intervals were calculated by bootstrap resampling from the data 1000 times with replacement. Datasets are available at GEO with accession number GSE153003.

### Fluorescence microscopy

Immunofluorescence of chromosome spreads was performed essentially as described (Rockmill, 2009). The antibodies used are listed in Table S3. Images of spreads and fixed whole cells were captured with a Nikon Eclipse 90i fluorescence microscope controlled with MetaMorph software and equipped with a Hamamatsu Orca-AG CCD camera and a PlanApo VC 100x 1.4 NA objective. The following exposure times were used: DAPI (400 msec), Mps3-GFP/Mps3-HA (500 msec), Cnm67-mCherry (200 msec), tubulin (10 msec) and H2A.Z-GFP (500 msec in wild type and 2000 msec in *swr1*).

For BiFC analysis, cells were fixed with 3.7% formaldehyde for 10 minutes at 30°C with 500 rpm shaking. Cells were washed with 1X PBS, permeabilized for 10 minutes with 70% ethanol and stained with 1 μg/ml DAPI for 10 minutes. Images were captured with the Nikon Eclipse 90i fluorescence microscope described above, with the following exposure times: DAPI (400 msec), Venus^YFP^ (5000 msec), Mps3-mCherry (1000 msec), Spc110-RedStar2 (1000 msec) and DIC (10 msec).

For analysis of Mps3-GFP distribution, cells were fixed with 3.7 % formaldehyde and washed with 1X PBS. Stacks of 30 planes at 0.2 μm intervals were captured for Mps3-GFP (300 msec exposure). Also, a DIC image (25 msec), and single-plane image of Hop1-mCherry (600 msec exposure), to identify meiotic prophase cells, were captured. Maximum intensity projections were generated using Fiji software (https://imagej.net/Fiji). Images were captured with an Olympus IX71 fluorescence microscope equipped with a personal DeltaVision system, a CoolSnap HQ2 (Photometrics) camera, and a 100x UPLSAPO 1.4 NA objective.

For colocalization of H2A.Z-GFP (400 msec exposure) and Mps3-MCherry (800 msec), Cnm67-mCherry (1000 msec) or Net1-RedStar2 (1000 msec) in live meiotic cells, z-stacks of 25 planes at 0.2 μm intervals were consecutively captured using the DeltaVision microscope described above. Images were deconvolved using the SoftWoRx 5.0 software (Applied Precisions).

For super-resolution analysis (SIM) of Mps3-GFP, H2A.Z-GFP and Spc110-mCherry, cells collected 16 h after meiotic induction were fixed for 15 min in 4% paraformaldehyde (Ted Pella) with 100 mM sucrose, and then washed two times in phosphate-buffered saline pH 7.4. Aliquots of cells were placed on cleaned slides covered with coverslips (number 1.5). Multiple color 3D-SIM images were acquired using a GE Healthcare DeltaVision OMX Blaze V3 fitted with an Olympus PlanApo N 100x 1.42 NA oil objective. Stacks of 17 planes at 0.125 μm intervals were captured (100 msec exposure for green and red channels). SIM reconstruction was performed with the Applied Precision SoftWoRx software package (GE Healthcare, Piscataway, NJ) following the Applied Precision protocols. After reconstruction, alignment between differently colored channels was performed based on calibration from alignment slide provided by the manufacturer. All analysis was performed using ImageJ and custom plugins written for ImageJ (created in the microscopy center of The Stowers Institute for Medical Research) at http://research.stowers.org/imagejplugins/index.html).

### Measurement of chromosome and telomere movement

For analysis of chromosome movement using Zip1-GFP tracking, cells from 16 h meiotic cultures of wild-type and mutants (*ndj1Δ, htz1Δ* or *swr1Δ*) were mixed in the same microscopy culture chamber (μ-slide 8 well, Ibidi) previously treated with 0.5 mg/ml of Concanavalin A Type IV (Sigma-Aldrich). The chamber was maintained at 30°C during the experiment. Zip1-GFP images were taken during 30 sec at 0.6 sec intervals with 100 msec exposure time. To distinguish wild-type cells (expressing *PMA1-mCherry*) from mutants, red channel images were also taken (800msec). Images were deconvolved using the SoftWoRx 5.0 software (Applied Precisions). Clearly isolated chromosomes in a nucleus were manually marked at the end and tracked for 50 consecutive frames. Chromosome velocities were calculated using a manual tracking plugin on ImageJ (https://imagej.nih.gov/ij/plugins/track/track.html). A total 16-25 chromosomes for each genotype in 4 independent experiments were analyzed.

For analysis of *TEL4L* movement, meiotic prophase cells from synchronous cultures (6 h after induction of *IME1* with CuSO4) were placed in Concanavalin A-treated microscopy culture chambers maintained at 30°C. For *TEL4L-GFP* (200 msec exposure) and Zip1-mCherry (800 msec exposure), Z-stacks of seven planes (0.6 μm step size) were captured at 8 sec intervals during 180 sec. A single plane of DIC was also captured in every frame. To correct for possible small displacements of the microscope stage during the time lapse, GFP images were aligned using DIC images as reference using a script provided by Giovanni Cardone (available upon request). *TEL4L-GFP* dots in the nuclei were manually marked and tracked for 23 consecutive frames. Telomere movement velocities were calculated using the MTrackJ plugin of Fiji (https://imagescience.org/meijering/software/mtrackj/). A total of 101-172 telomere tracks from 3 different experiments were analyzed for each genotype. Images for both chromosome movement (Zip1-GFP) and telomere movement (*TEL4L-GFP*) were captured with an Olympus IX71 fluorescence microscope equipped with a personal DeltaVision system, a CoolSnap HQ2 (Photometrics) camera, and a 100x UPLSAPO 1.4 NA objective.

### Statistics

To determine the statistical significance of differences a two-tailed Student *t*-test, for pairwise comparisons, or one-way ANOVA Tukey test, for multiple comparisons, were used. *P*-Values were calculated with the GraphPad Prism 5.0 software. The nature of errors bars in graphical representations and the number of biological replicates is indicated in the corresponding figure legend.

## ACKNOWLEDGMENTS

We thank Andrés Clemente and Félix Prado for useful comments and discussion, Cristina Martín, Andrés Clemente, César Roncero, Raimundo Freire and Shirleen Roeder for reagents, and Giovanni Cardone for the image alignment script. This work was supported by a grant BFU2015-65417-R from Ministry of Economy and Competitiveness (MINECO) of Spain to PSS, a grant RTI2018-099055-B-I00 from Ministry of Science, Innovation and Universities (MCIU/AEI/FEDER, UE) of Spain to PSS and JAC, the Stowers Institute for Medical Research to SLJ, a grant R01GM121443 from the National Institute of General Medical Sciences of the National Institutes of Health to SLJ, and NIH grants GM111715 and GM123035 to AH. SG-A was partially supported by the “Unidad de Excelencia de Biología Funcional y Genómica” Ref. 2019/X002/05 from the University of Salamanca. The IBFG is supported in part by an institutional grant from the “Junta de Castilla y León, Ref. CLU-2017-03 co-funded by the P.O. FEDER de Castilla y León 14-20”. Original data underlying parts of this manuscript can be accessed from the Stowers Original Data Repository at http://www.stowers.org/research/publications/LIBPB-1519.

## AUTHOR CONTRIBUTIONS

S González-Arranz: Conceptualization, investigation, formal analysis, visualization

JM Gardner: Investigation

Z Yu: Formal analysis

NJ Patel: Investigation, formal analysis

J Heldrich: Investigation, formal analysis

B Santos: Supervision, project administration

JA Carballo: Resources, funding acquisition

SL Jaspersen: Resources, funding acquisition

A Hochwagen: Resources, formal analysis, funding acquisition

PA San-Segundo: Conceptualization, investigation, supervision, funding acquisition, visualization, writing original draft

All authors revised, commented and approved the manuscript.

## Non-standard abbreviations

BiFC: Bimolecular Fluorescence Complementation
DIC: Differential Interference Contrast
KAc: Potassium Acetate
NE: Nuclear Envelope
LINC: Linker of the Nucleoskeleton and Cytoskeleton
rDNA: Ribosomal DNA
SC: Synaptonemal Complex
SIM: Structured Illumination Microscopy
SPB: Spindle Pole Body
VC: C-terminal moiety of the Venus fluorescent protein
VN: N-terminal moiety of the Venus fluorescent protein
WCE: Whole cell extracts

---

**Figure S1.**
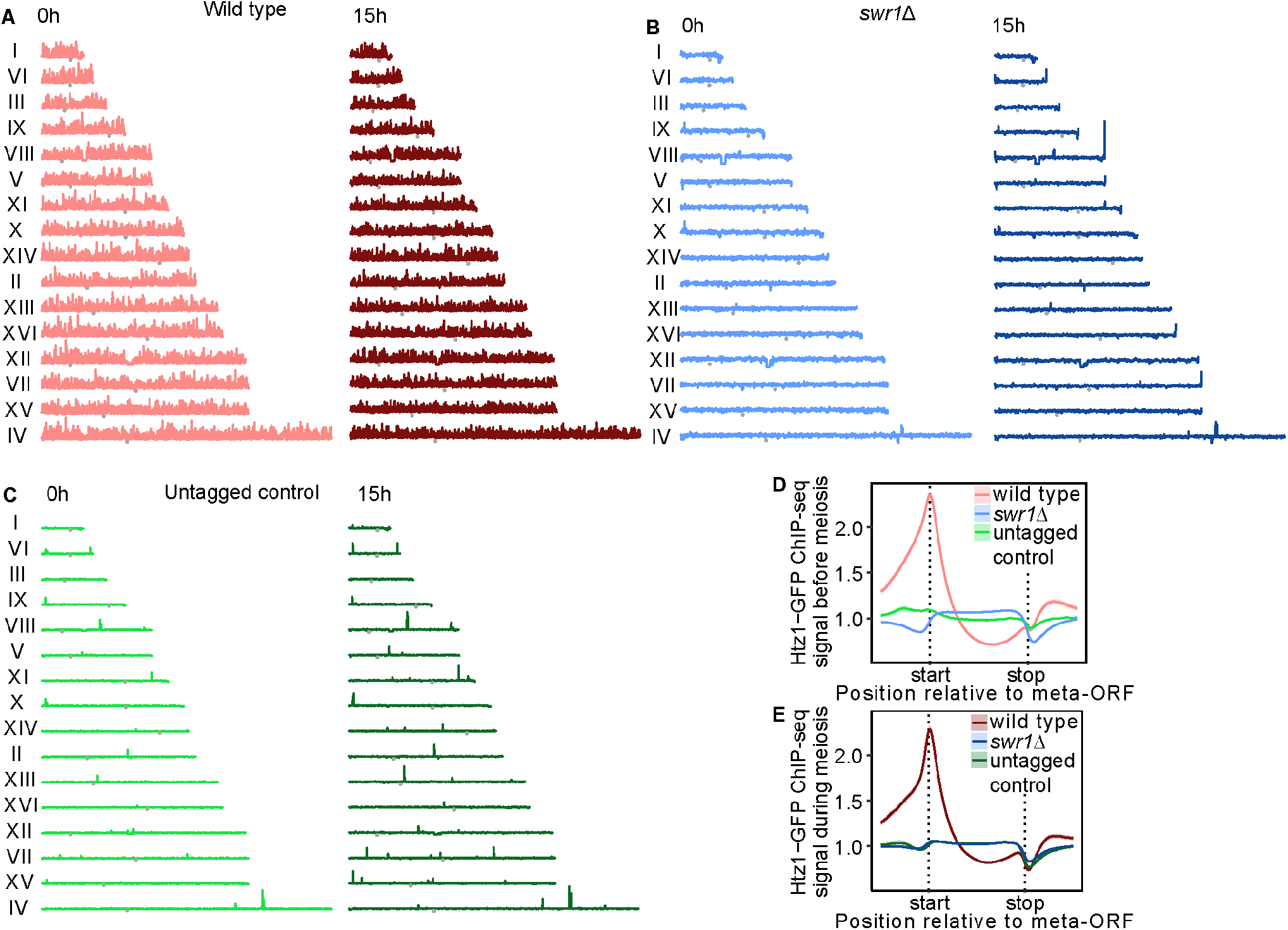
Genome-wide incorporation of H2A.Z to meiotic chromatin depends on SWR1. (Related to Figure 1) Profiles of H2A.Z binding to all chromosomes in wild type **(A)**, *swr1Δ* **(B)**, and the untagged control **(C)**, determined by ChIP-seq. Gray points indicate the location of the centromere. **(D-E)** Metagene analysis of H2A.Z binding by ChIP-seq. The ORFs are scaled to the “Start” and “Stop” positions, and up- and downstream flanking regions represent half the size of the ORF. Samples were taken at 0 h and 15 h after meiotic induction. Anti-GFP antibodies were used to immunoprecipitate H2A.Z-GFP. Strains are: DP840 (*HTZ1-GFP*), DP841 (*HTZ1-GFP swr1Δ*) and DP421 (*HTZ1* untagged control).

**Figure S2.**
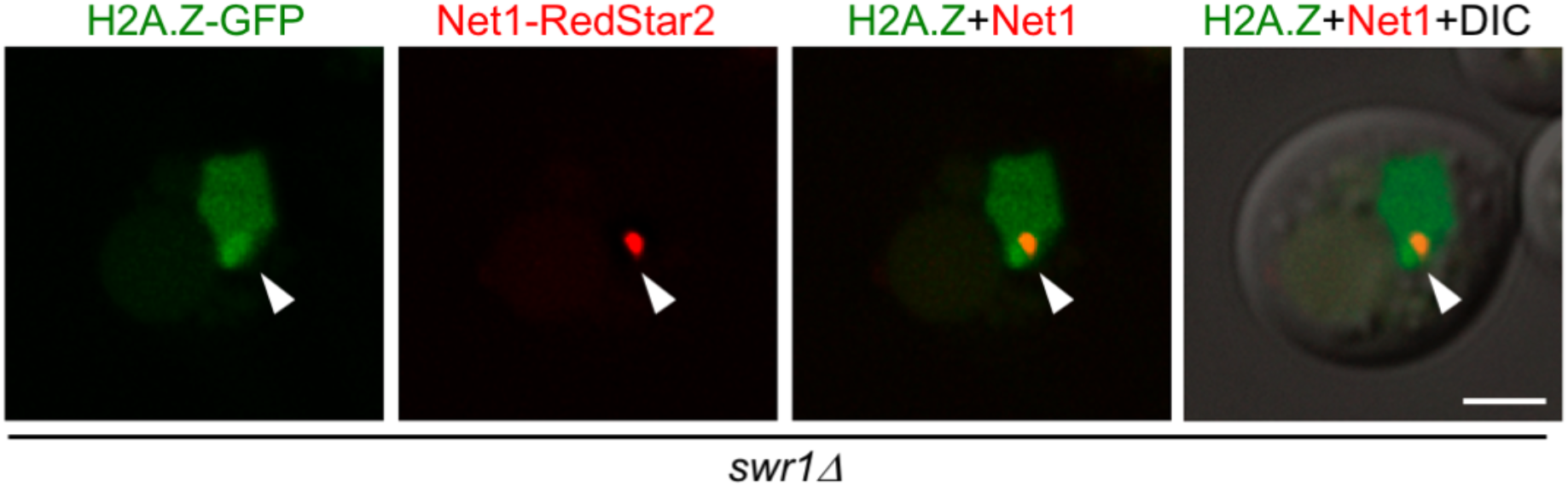
A fraction of H2A.Z accumulates in the vicinity of the nucleolus in *swr1Δ* (Related to Figure 2) Microscopy fluorescence images of *swr1Δ* cells expressing *HTZ1-GFP* and *NET1-RedStar2* as a nucleolar marker. A single plane of a representative cell displaying a diffuse peripheral accumulation of H2A.Z is shown. The arrowhead points to the nucleolar area marked by Net1. Images were taken 16 h after meiotic induction. Scale bar, 2 μm. The strain is DP1189 (*swr1Δ HTZ1-GFP NET1-RedStar2*)

**Figure S3.**
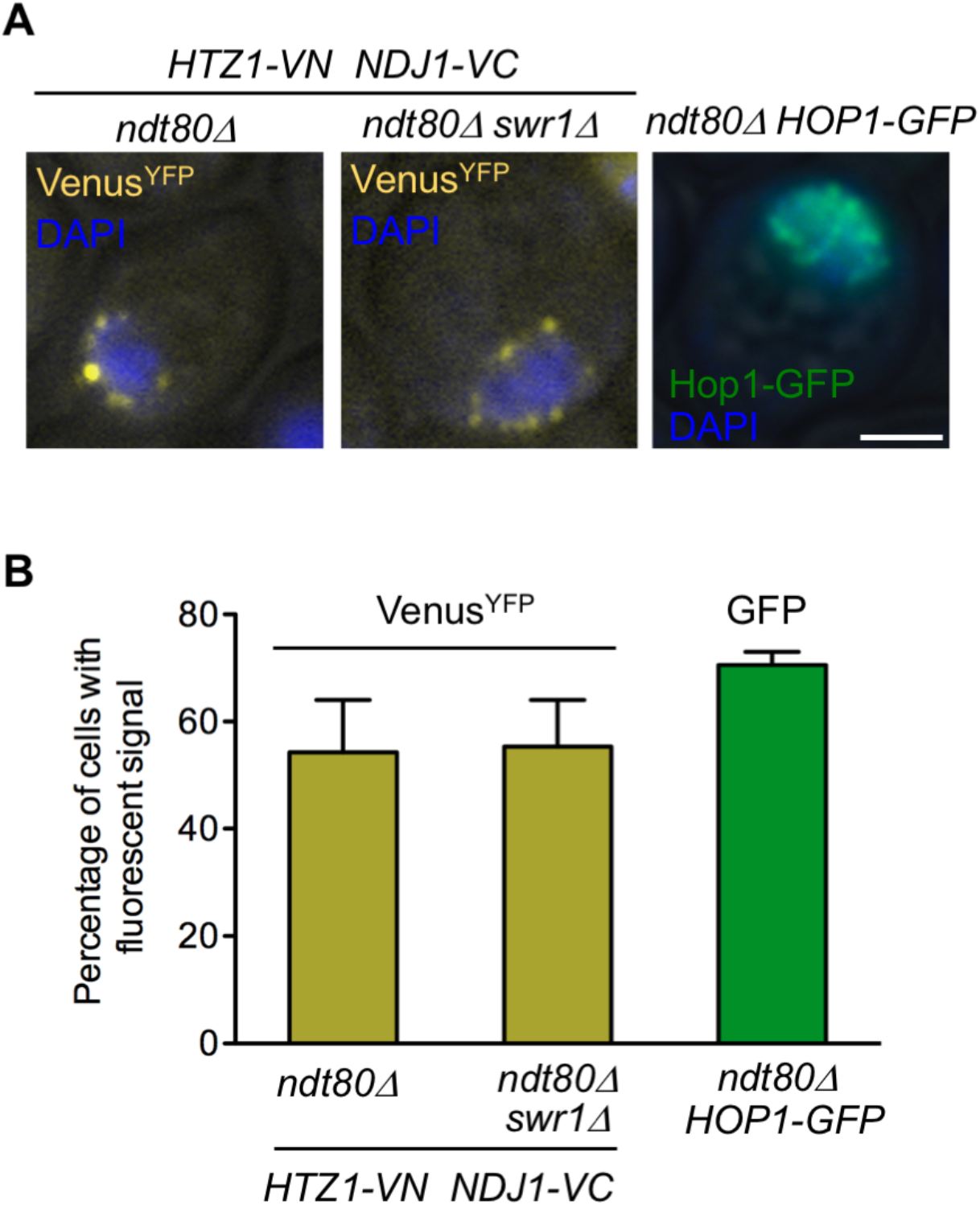
BiFC analysis of H2A.Z-Ndj1 interaction in *ndt80Δ* cells (Related to Figure 3). **(A)** Microscopy fluorescence images of *ndt80Δ* and *ndt80Δ swr1Δ* cells expressing *HTZ1* fused to the N-terminal half of the Venus^YFP^ (*VN*) and *NDJ1* fused to the C-terminal half of the Venus^YFP^ (*VC*). Nuclei are stained with DAPI (blue). The reconstitution of Venus^YFP^ fluorescence resulting from H2A.Z-VN/Ndj1-VC interaction appears in yellow. A parallel meiotic culture of *ndt80Δ* cells expressing *HOP1-GFP* (green) was used as control for meiotic prophase I staging. Images were taken 24 h after meiotic induction. Representative cells are shown. Scale bar, 2 μm **(B)** Quantification of the percentage of cells displaying Venus^YFP^ fluorescent signal or Hop1-GFP signal, as indicated. The analysis was performed in triplicate. More than 300 cells were scored in every experiment. Error bars, SD. Strains are: DP1748 (*ndt80Δ HTZ1-VN NDJ1-VC*), DP1749 (*ndt80Δ swr1Δ HTZ1-VN NDJ1-VC*) and DP963 (*ndt80Δ HOP1-GFP*).

**Figure S4.**
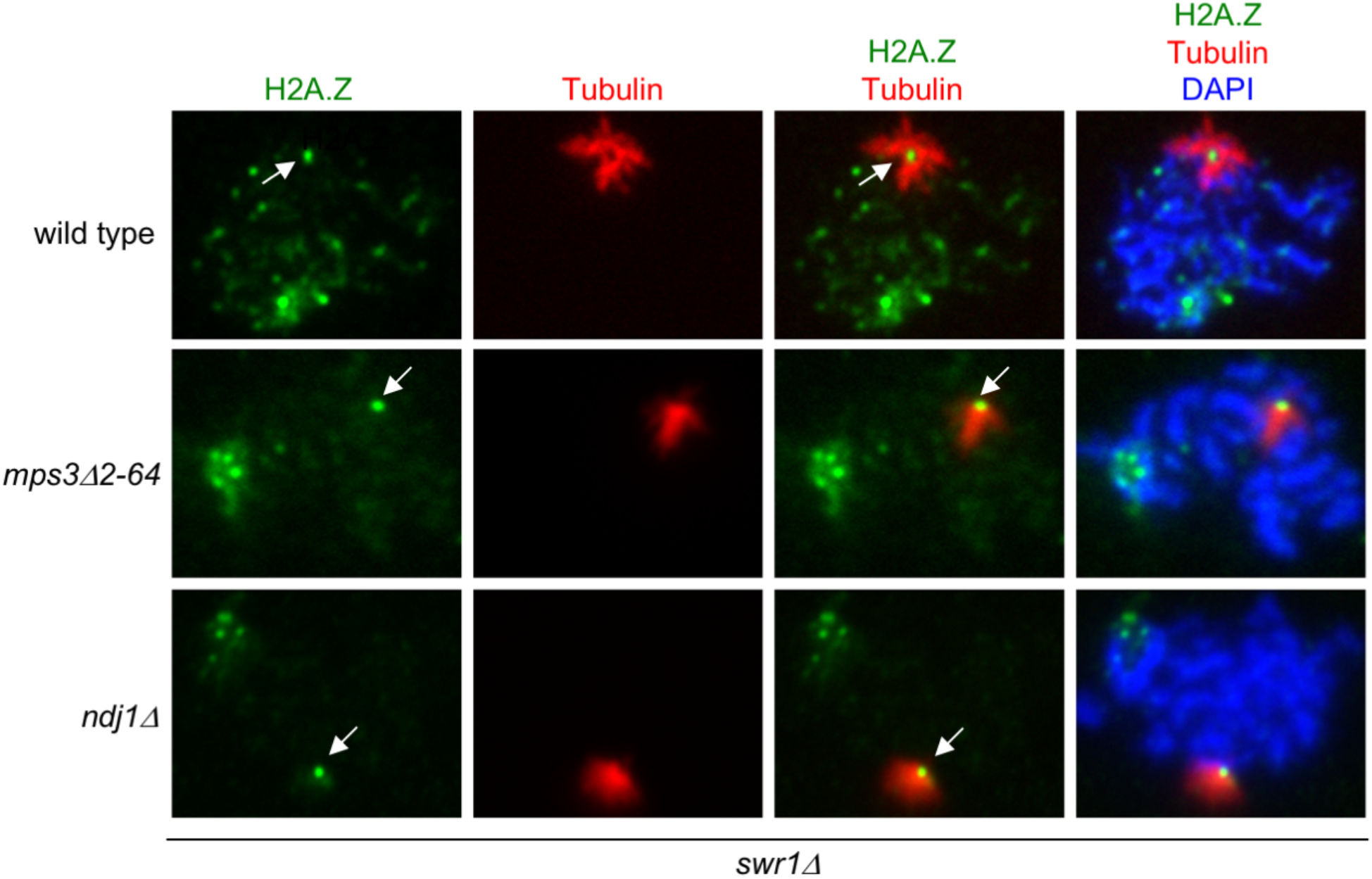
Localization of H2A.Z to the SPB is independent of Ndj1 and the 2-64 N-terminal domain of Mps3. (Related to Figure 5) Immunofluorescence of representative spread pachytene nuclei stained with DAPI to visualize chromatin (blue), anti-GFP to detect H2A.Z (green), and anti-tubulin to mark the monopolar prophase spindle (red). The arrow points to an H2A.Z focus present at the center of the bushy spindle corresponding to the SPB location. Strains are DP1395 (wild type), DP1280 (*mps3-Δ2-64*) and DP1305 (*ndj1Δ*). 25, 21 and 23 nuclei were examined for wild type, *mps3-Δ2-64* and *ndj1Δ*, respectively.

**Figure S5.**
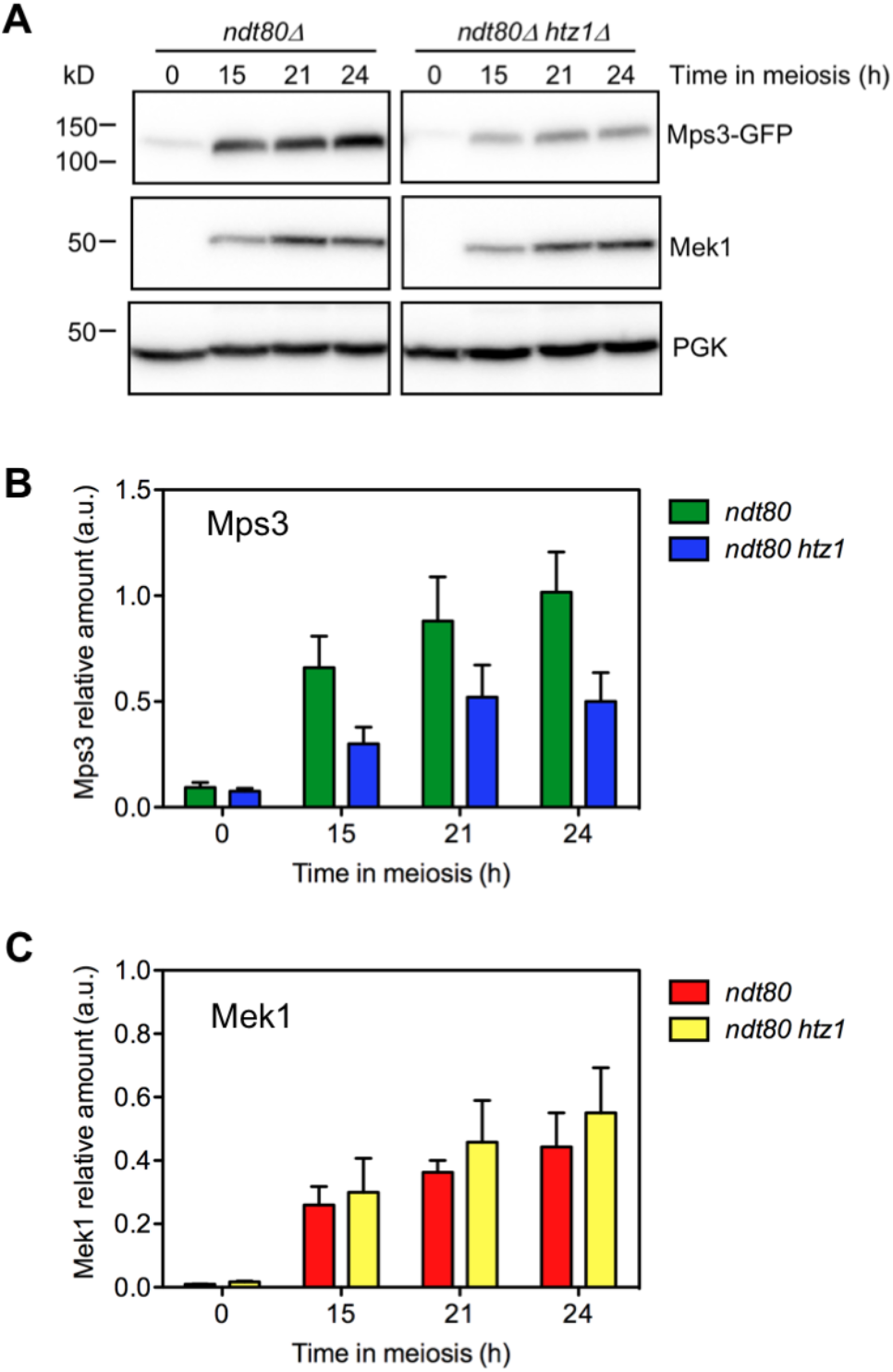
Mps3 global levels are reduced in *ndt80Δ-arrested* cells lacking H2A.Z. (Related to Figure 6) **(A)** Western blot analysis of Mps3-GFP and Mek1 production during meiosis detected with anti-GFP and anti-Mek1 antibodies, respectively. PGK was used as a loading control. A representative blot is shown. **(B-C)** Quantification of Mps3-GFP (B) and Mek1 (C) levels normalized to PGK. Average and SEM (error bars) from three independent experiments are shown. Strains are: DP1014 (*ndt80Δ MPS3-GFP*) and DP1013 (*ndt80Δ htz1Δ MPS3-GFP*).

**Figure S6.**
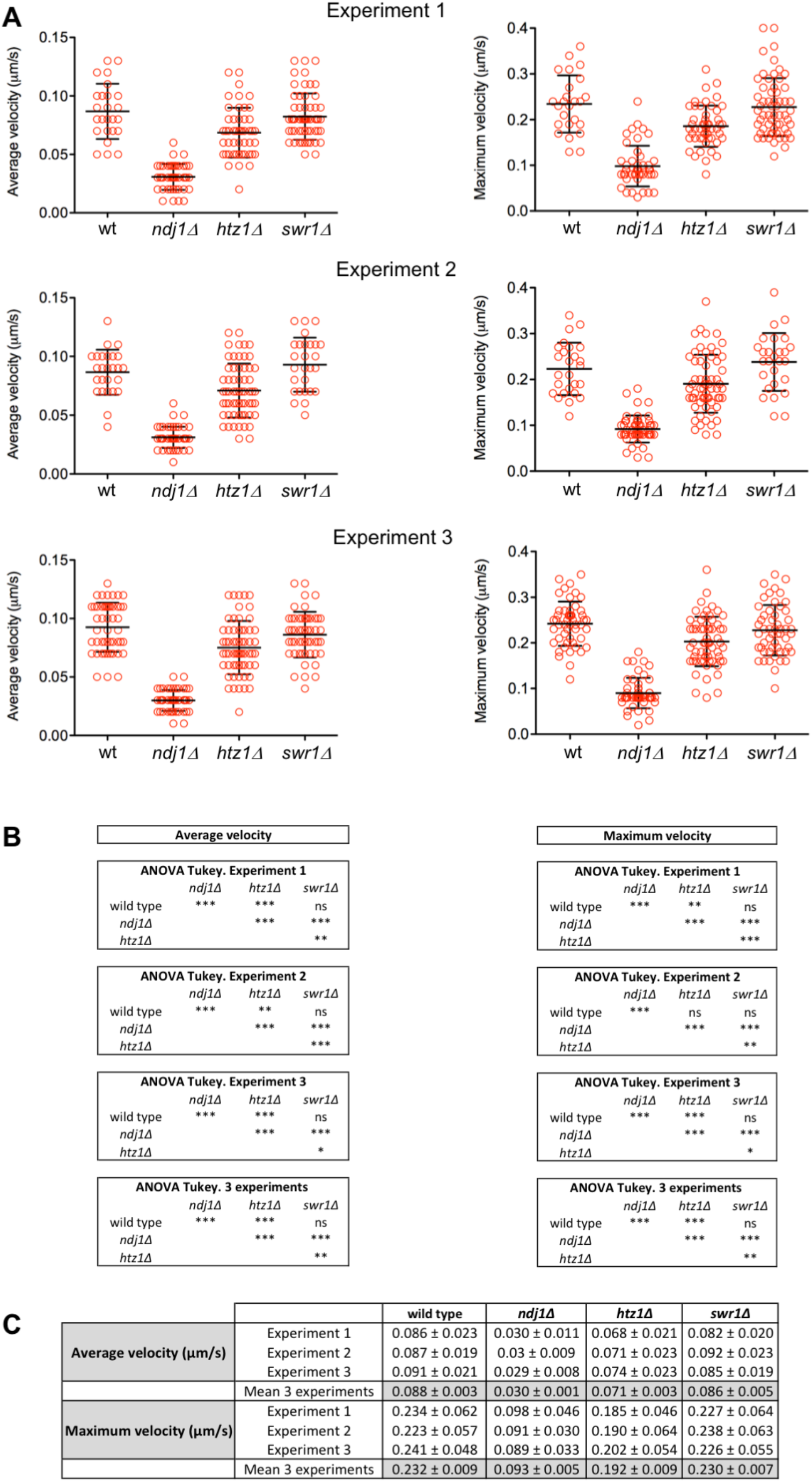
Analysis of *TEL4L* movement. (Related to Figure 8). **(A)** Measurement of average velocity and maximum velocity in three independent time-lapse experiments tracking *TEL4L* movement marked with GFP as depicted in Figure 8. Error bars, SD. **(B)** ANOVA statistical analysis of the velocity data obtained in every individual experiment as well as combining the data from all three experiments. **(C)** Mean values for average and maximum velocity. Strains are DP1692 (wild type), DP1722 (*ndj1Δ*), DP1693 (*htz1Δ*) and DP1694 (*swr1Δ*).

**Table S1.** *Saccharomyces cerevisiae* strains.

| Strain | Genotype | Source |
| --- | --- | --- |
| DP421 | *MATa/MATα leu2,3-112 his4-260 thr1-4 ura3-1 trp1-289 ade2-1 lys2ΔNheI* | PSS Lab |
| DP838 | DP421 *htz1::URA3 ZIP1-GFP* | This work |
| DP840 | DP421 *HTZ1-GFP::kanMX6* | PSS Lab |
| DP841 | DP421 *swr1::natMX4 HTZ1-GFP::kanMX6* | PSS Lab |
| DP866 | DP421 *MPS3-GFP::kanMX6* | This work |
| DP867 | DP421 *htz1::URA3 MPS3-GFP::kanMX6* | This work |
| DP957 | DP421 *ndj1::natMX4 ZIP1-GFP* | This work |
| DP963 | DP421 *ndt80::LEU2 HOP1-GFP::kanMX6* | PSS Lab |
| DP1013 | DP421 *ndt80::LEU2 htz1::URA3 MPS3-GFP::kanMX6* | This work |
| DP1014 | DP421 *ndt80::LEU2 MPS3-GFP::kanMX6* | This work |
| DP1032 | DP421 *MPS3-GFP::kanMX6 HOP1-mCherry::natMX4* | This work |
| DP1033 | DP421 *htz1::URA3 MPS3-GFP::kanMX6 HOP1-mCherry::natMX4* | This work |
| DP1057 | DP421 *ZIP1-GFP PMA1-mCherry::natMX4* | This work |
| DP1091 | DP421 *swr1::natMX4 ZIP1-GFP* | This work |
| DP1102 | DP421 *swr1::natMX4 MPS3-GFP::kanMX6* | This work |
| DP1103 | DP421 *ndj1::natMX4 MPS3-GFP::kanMX6* | This work |
| DP1108 | DP421 *swr1::natMX4 HTZ1-GFP::kanMX6 MPS3-mCherry::hphMX4* | This work |
| DP1172 | DP421 *HTZ1-GFP::kanMX6 CNM67-mCherry::natMX4 swr1::hphMX4* | This work |
| DP1182 | DP421 *HTZ1-GFP::kanMX6 swr1::hphMX4* | This work |
| DP1189 | DP421 *HTZ1-GFP::kanMX6 NET1-RedStar2::natNT2 swr1::hphMX4* | This work |
| DP1280 | DP421 *mps3::hphMX4 pSS326 [mps3Δ2-64-mCherry URA3] HTZ1-GFP::kanMX6 swr1::natMX4* | This work |
| DP1305 | DP421 *HTZ1-GFP::kanMX6 swr1::natMX4 ndj1::hphMX4* | This work |
| DP1330 | DP421 *MPS3-3HA::kanMX6* | This work |
| DP1394 | DP421 *MPS3-3HA::natMX4 HTZ1-GFP::kanMX6* | This work |
| DP1395 | DP421 *swr1::hphMX4 MPS3-3HA::natMX4 HTZ1-GFP::kanMX6* | This work |
| DP1493 | DP421 *swr1::hphMX4 HTZ1-VN::TRP1 NDJ1-VC::kanMX6* | This work |
| DP1496 | DP421 *HTZ1-VN::TRP1 NDJ1-VC::kanMX6* | This work |
| DP1506 | DP421 *SPC110-RedStar2::natNT2/SPC110 HTZ1-VN::TRP1 NDJ1-VC::kanMX6 swr1::hphMX4* | This work |
| DP1511 | DP421 *mps3::natMX4 pSS269 [MPS3-mCherry URA3] HTZ1-VN::TRP1 NDJ1-VC::kanMX6* | This work |
| DP1512 | DP421 *mps3::natMX4 pSS326 [mps3Δ2-64-mCherry URA3] HTZ1-VN::TRP1 NDJ1-VC::kanMX6* | This work |
| DP1540 | DP421 *HTZ1-VN::TRP1* | This work |
| DP1541 | DP421 *NDJ1-VC::kanMX6* | This work |
| DP1576 | DP421 *MPS3-GFP::kanMX6 swr1::natMX4 SPC110-mCherry::hphNT1/SPC110* | This work |
| DP1578 | DP421 *HTZ1-GFP::kanMX6 swr1::natMX4 SPC110-mCherry::hphNT1/SPC110* | This work |
| DP1692 | DP421 *P<sub>CUP1</sub>-IME1::kanMX6 ZIP1-mCherry/ZIP1 TEL4L-tetO(50)::URA3/TEL4L TetR-GFP::LEU2/leu2* | This work |
| DP1693 | DP421 *htz1::natMX4 P<sub>CUP1</sub>-IME1::kanMX6 ZIP1-mCherry/ZIP1 TEL4L-tetO(50)::URA3/TEL4L TetR-GFP::LEU2/leu2* | This work |
| DP1694 | DP421 *swr1::hphMX4 P<sub>CUP1</sub>-IME1::kanMX6 ZIP1-mCherry/ZIP1 TEL4L-tetO(50)::URA3/TEL4L TetR-GFP::LEU2/leu2* | This work |
| DP1722 | DP421 *ndj1::kanMX6 P<sub>CUP1</sub>-IME1::kanMX6 ZIP1-mCherry/ZIP1 TEL4L-tetO(50)::URA3/TEL4L TetR-GFP::LEU2/leu2* | This work |
| DP1748 | DP421 *ndt80::natMX4 HTZ1-VN::TRP1 NDJ1-VC::kanMX6* | This work |
| DP1749 | DP421 *ndt80::natMX4 swr1::hphMX4 HTZ1-VN::TRP1 NDJ1-VC::kanMX6* | This work |

\*All strains are diploids isogenic to BR1919 (Rockmill and Roeder, 1990). Unless specified, all strains are homozygous for the indicated markers. DP421 is a *lys2* version of the original BR1919-2N.
Rockmill, B., and G.S. Roeder. 1990. Meiosis in asynaptic yeast. *Genetics*. 126:563-574.

**Table S2.**
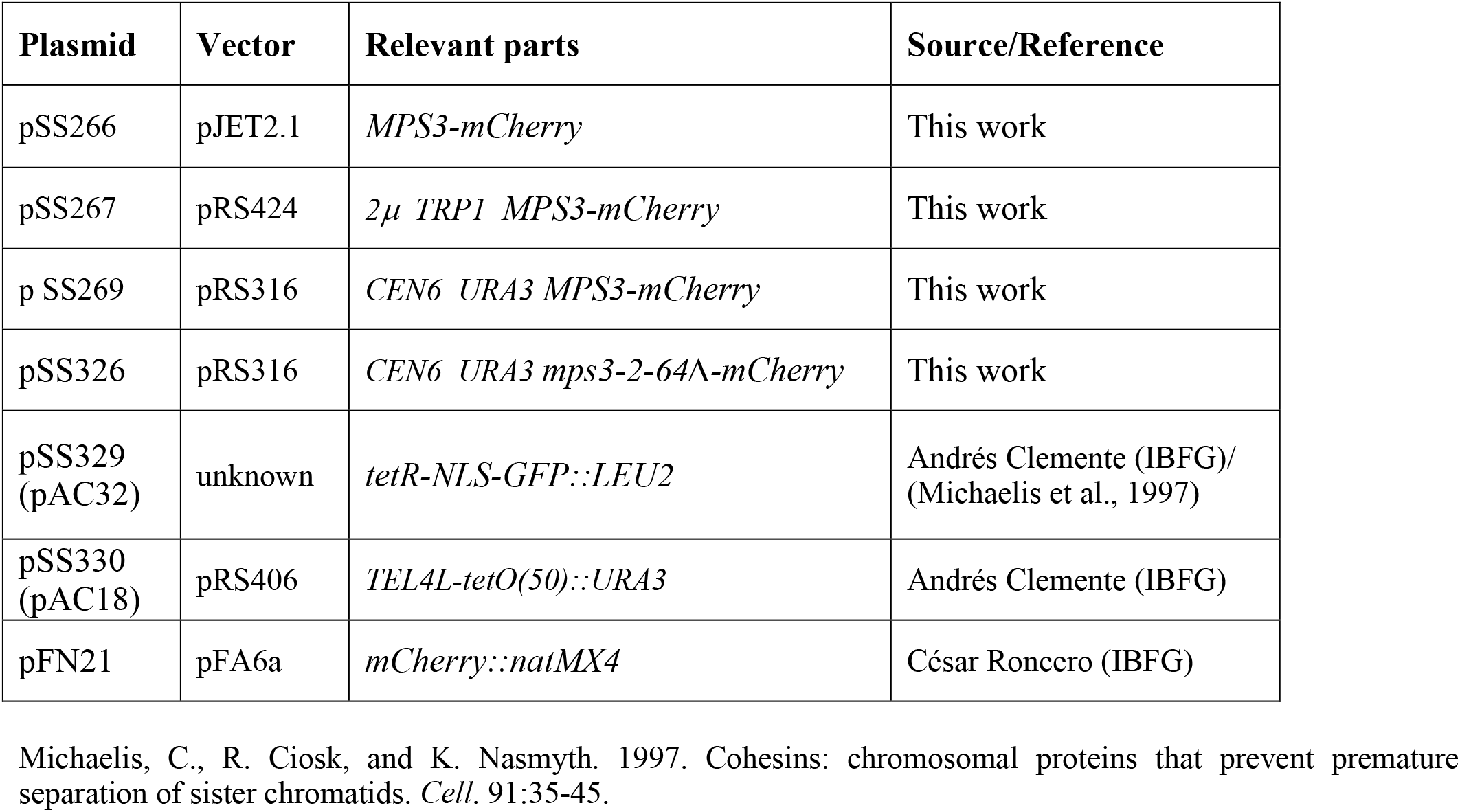
Plasmids.

| Plasmid | Vector | Relevant parts | Source/Reference |
| --- | --- | --- | --- |
| pSS266 | pJET2.1 | *MPS3-mCherry* | This work |
| pSS267 | pRS424 | *2μ TRP1 MPS3-mCherry* | This work |
| p SS269 | pRS316 | *CEN6 URA3 MPS3-mCherry* | This work |
| pSS326 | pRS316 | *CEN6 URA3 mps3-2-64Δ-mCherry* | This work |
| pSS329<br>(pAC32) | unknown | *tetR-NLS-GFP::LEU2* | Andrés Clemente (IBFG)/<br>(Michaelis et al., 1997) |
| pSS330<br>(pAC18) | pRS406 | *TEL4L-tetO(50)::URA3* | Andrés Clemente (IBFG) |
| pFN21 | pFA6a | *mCherry::natMX4* | César Roncero (IBFG) |

Michaelis, C., R. Ciosk, and K. Nasmyth. 1997. Cohesins: chromosomal proteins that prevent premature separation of sister chromatids. *Cell*. 91:35-45.

**Table S3.** Primary and secondary antibodies.

| **Antibody** | **Host and type** | **Application\***<br>**(Dilution)** | **Source / Reference** |
| --- | --- | --- | --- |
| H2A (acidic patch) | Rabbit polyclonal | WB (1:5000) | Merck 07-146 |
| H2A.Z | Rabbit polyclonal | WB (1:1000) | Active Motif 39647 |
| H2B | Rabbit polyclonal | WB (1:5000) | Abcam ab1790 |
| H3 | Rabbit polyclonal | WB (1:5000) | Abcam ab1791 |
| H4 | Rabbit polyclonal | WB (1:1000) | Abcam ab10158 |
| Zip1 | Rabbit polyclonal | IF (1:300) | S. Roeder |
| HA (12CA5) | Mouse monoclonal | WB (1:1000) | Roche 11666606001 |
| HA (3F10) | Rat monoclonal | IF (1:225) | Roche 11867431001 |
| GFP (JL-8) | Mouse monoclonal | WB (1:1000-1:2000)<br>IF (1:200) | Clontech 632381 |
| GFP | Rabbit polyclonal | ChIP-seq | R. Freire |
| mCherry/DsRed | Rabbit polyclonal | IF (1:200) | Clontech 632496 |
| Pgk1 (22C5D8) | Mouse monoclonal | WB (1:5000) | Invitrogen 459250 |
| Mek1 | Rabbit polyclonal | WB (1:1000) | Ontoso et al., 2013 |
| Tubulin | Rabbit monoclonal | IF (1:500) | Abcam EPR13798 |
| Anti-mouse-HRP | Sheep polyclonal | WB (1:5000) | GE-Healthcare NA931 |
| Anti-rabbit-HRP | Donkey polyclonal | WB (1:5000) | GE-Healthcare NA934 |
| Anti-mouse AF488 | Goat polyclonal | IF (1:200) | Invitrogen A11029 |
| Anti-rabbit AF594 | Goat polyclonal | IF (1:200) | Invitrogen A11012 |
| Anti-rat AF568 | Goat polyclonal | IF (1:200) | Invitrogen A11077 |

\*WB, western blot; IF, immunofluorescence; ChIP-seq, chromatin immunoprecipitation-sequencing
Ontoso, D., I. Acosta, F. van Leeuwen, R. Freire, and P.A. San-Segundo. 2013. Dot1-dependent histone H3K79 methylation promotes activation of the Mek1 meiotic checkpoint effector kinase by regulating the Hop1 adaptor. *PLoS Genet.* 9:e1003262.

